# Spike protein E2 of chikungunya virus: a plant-based vaccine exhibited potent immunogenicity in BALB/c mice

**DOI:** 10.1101/2025.08.21.671458

**Authors:** Sadia Qamar, Monika Dalal, Amna Syeda, Malik Zainul Abdin, Firdaus Qamar, Altaf Ahmad, Md. Amjad Beg, S. Naved Quadri, Javed Ahmad, Suhel Parvez, M. Irfan Qureshi

**Affiliations:** Department of Biotechnology, Faculty of Life Sciences, Jamia Millia Islamia, New Delhi - 110025, India; Department of Biotechnology, ICAR-National Institute for Plant Biotechnology, IARI, New Delhi - 110012, India; Department of Biotechnology, School of Chemical and Life Sciences, Jamia Hamdard, New Delhi, India; Department of Botany, Aligarh Muslim University, Aligarh - 202002, India; Rutgers Cancer Institute of New Jersey, The State University of New Jersey, New Brunswick, NJ, USA; Oak Ridge National Laboratory, Oak Ridge, TN 37831, United States; Department of Toxicology, School of Chemical and Life Sciences, Jamia Hamdard, New Delhi - 110062, India

**Keywords:** Chikungunya fever, recombinant E2 protein, plant-based vaccine, active immunity

## Abstract

Over the last two decades, chikungunya virus (CHIKV) infections have surged worldwide, causing significant suffering. CHIKV E2 gene codes for spike protein E2, essential for virus-host interactions and thus serves as a potential vaccine candidate. We expressed a full-length E2 (S27 African prototype) in both *E. coli* and *Nicotiana tabacum*, which triggered an immune response in BALB/c mice. First, E2 was computationally analyzed for PTM patterns and prediction of B-cell and T-cell epitopes. Next, molecular docking of epitopes with MHCs (Class I and II) revealed high affinity, confirmed by Molecular Dynamics Simulation. Then, the chemically synthesized E2-6xHis tag was cloned and expressed in both *E. coli* and *N. tabacum* under T7 and CaMV promoters, respectively. E2 was cloned into the pUC57 vector (*E. coli*) and expressed using the pET28a(+) vector in BL21(DE3)pLysS cells, followed by cloning into the pCAMBIA1302 vector and transformation into *Agrobacterium tumefaciens*. RT-PCR and confocal visualization confirmed the formation of E2 transcripts. Recombinant E2 was purified on Ni-NTA columns and visualized as a protein of ∼49 kDa on SDS PAGE. Finally, E2 was injected into BALB/c mice; neutralizing antibodies, including IgG, were detected as positive in the indirect ELISA, with the highest levels observed at 3 days post-infiltration (3 dpi). Western blot also confirmed E2 expression in *E coli* and tobacco, and induction of E2-specific antibodies in BALB/c mice. This study presents a promising approach to developing a safe and effective vaccine against chikungunya fever in plants.

## INTRODUCTION

Chikungunya virus (CHIKV) is a mosquito-borne virus transmitted primarily by *Aedes* mosquitoes. It causes acute illness with body rashes, fever, debilitating joint pain, and a curved posture known as “Makonde”. Many patients experience long-term disabilities and joint pain. Chikungunya fever is a significant health concern that accounts for a huge health and economic burden. In June 2025, the WHO made an emergency update through the Epidemic and Pandemic Preparedness and Prevention (EPP) team with a warning of the potential re-emergence in previously affected areas. So far, 119 countries and territories have documented indigenous mosquito-borne transmission of CHIKV (Weber et al., 2024; WHO Global Chikungunya Epidemiology Update – June 2025). Climate change is further held responsible for the widespread of disease because it favours mosquito breeding (Romanello et al. 2021). Therefore, Africa, the Indian subcontinent, and Southeast Asia often see sporadic outbreaks of Chikungunya virus infection in large numbers (Russo et al. 2020). The Chikungunya virus was discovered in 1953 in Newala, Tanzania, and initially thought to be related to dengue. CHIKV belongs to the Togaviridae family, Alphavirus genus, and shares ancestry with the O’nyong-nyong virus (ONNV) (Kendall et al., 2019; Powers et al., 2000). CHIKV, a spherical virus with a 70 nm diameter, has an 11.8 kb single-stranded positive RNA genome. The genome has two untranslated regions, indicating adaptation to its mosquito vector. It contains two open reading frames (ORFs) encoding precursor proteins. The 5’ ORF produces non-structural proteins forming the viral replication complex, while the 3’ ORF produces structural proteins (Metz and Pijlman, 2016).

Chikungunya virus enters human cells through receptor-mediated endocytosis and pH-dependent fusion, common to other alphaviruses. Surface proteins E2 and E1 mediate cell entrance, forming trimers of heterodimers. E2 protein domains A, B, and C, with N-glycosylation sites, are potential sites for target cell contact. The structural protein E1 helps in membrane fusion (Li et al., 2010), while the E2 protein is responsible for receptor binding (Fox et al., 2015). As a result, E1 and E2 proteins are good candidates for vaccinations as immunogens. Numerous investigations have demonstrated the significance and immunodominance of structural protein E2. Throughout the infection, E2 is the primary target of specific antibodies in infected patients (Kam et al., 2012a, 2012b). Importantly, E2-neutralizing antibodies produced in animals can prevent the virus from fusing, binding, or entering the target cell (Smith et al., 2015). Furthermore, anti-E2 antibodies from recovering patients can be passively transferred to adults and newborns to minimize or completely eradicate CHIKV infection (Fox et al., 2015). Moreover, structural envelope proteins serve as the foundation for the majority of CHIKV subunit vaccine candidates that have advanced to clinical trials (Reyes-Sandoval, 2019).

Until November 9, 2023, no licensed antiviral vaccine existed. Ixchiq, the first FDA-approved vaccine (Weber et al., 2024), has limitations and is causing side effects due to its live, weakened Chikungunya virus content, as mentioned in Phase 3 trials of VLA1553. Another vaccine, which completed a trial in September 2020, PXVX0317 (CHIKV VLP), showed many adverse events (Bennett et al., 2022).

There are no reports on cloning and expression of full-length S27-African prototype E2 gene/protein in plants and subsequent immunogenic studies in the BALB/c mouse model. Therefore, given the importance of the E2 protein as a potent vaccine target against CHIKV, the present study describes the cloning and expression of the E2 gene from the S27-African strain in *E. coli* and *N. tabacum*, and demonstrates an immunogenic response in BALB/c mice. The study is unique in its development of the CHIKV vaccine in plant systems.

## Materials and Methods

### Phylogenetic analysis

The E2 protein of the Chikungunya virus sequence was searched for significant alignment using the BLASTp suite in the phylogeny reconstruction map. A neighbor-joining tree was constructed using MEGA12 (Kumar et al., 2024). A focused phylogenetic tree (0.01 distance relation and bootstrap 100) serotype was generated using the Chikumgunya Typing Tool, Genome Detective Platform ().

### Comprehensive structural analysis

CHIKV E2 was analysed for the frequency of amino acid residues and their significance. Possible post-translational modifications that rE2 may undergo in humans were predicted using the “sitetack” portal (https://sitetack.net) (Gutierrez et al., 2024). Parameters analyzed included the frequency of different amino acids and the probability prediction of post-translational modification in the human body. Computational prediction for possible PTM sites was performed and compared against the human proteome with actual PTMs using PTMGPT2 portal (Shrestha et al., 2024). PTMs targeted were Amidation (V), Methylation (K), Ubiquitination (K), Phosphorylation (S,T), N-linked Glycosylation (N), S-nitrosylation (C), Phosphorylation (Y), Acetylation (K), Methylation (R), Hydroxylation (K), Hydroxylation (P), O-linked Glycosylation (S,T), Malonylation (K), Glutarylation (K), Glutathionylation (C), S-palmitoylation (C), Succinylation (K), Sumoylation (K), and Formylation (K). Similarly, the MusiteDeep (https://www.musite.net) (Wang et al., 2020) was used with 0.5 threshold for general prediction of PTM-able residues considering that this will be true for plants including tobacco where recombinant protein was expressed. Validation of data was performed using VPTMdb, a viral PTM database (Xiang et al., 2021) taking top ten E2 sequences, including Q8JUX5 as the model protein sequence.

### Prediction of B-cell epitopes

The gene (GeneBank Accession No. AF339485.1; Supplementary Table 1) and protein sequence (ID: Q8JUX5; starting “STKDNFNVYK”, 47.88 kDa; Supplementary Table 2) were taken into consideration in our study. Various tools used for the prediction of B cell epitopes were Bepipred linear Epitope prediction (Larsen et al., 2006), Parker Hydrophilicity (Parker et al., 1986), Emini Surface Accessibility (Emini et al., 1985), Chou and Fasman Beta Turn (Chou et al., 1978), Kolaskar and Tongaonkar Antigenicity (Kolaskar et al., 1990), and Karplus and Schulz flexibility prediction (Karplus et al., 1985). The jobs on the said servers were run between 15 to 25 February 2023.

### Prediction of T-cell epitopes

The Chikungunya virus Structural Polyprotein (SP) sequence (ID: Q8JUX5) was acquired from the UniProt database in FASTA format. Employing the NETCTL_1.2 server with a threshold value of 0.95 (ensuring sensitivity and specificity of 0.90 and 0.95), CD8+ T cell peptides were identified. Utilizing 12 MHC Class-I supertypes, including A1, A2, A3, A24, A26, B7, B8, B27, B39, B44, B58, and B62, we predicted epitopes based on the combined score of C-terminal cleavage and TAP transport efficiency. Peptides with the highest overall combined score were chosen as epitope candidates. TepiTool (iedb.org) was then used to assess peptide binding to MHC Class-I molecules, with predicted outputs in IC50 nM units (ranging from < 50 nM to < 500 nM), indicating high to moderate affinity.

The MHC-NP tool, employing machine learning, accurately predicted peptide binding to specific MHC molecules. The Class I Immunogenicity tool was employed to predict the immunogenicity of MHC Class I binding peptides. Additionally, the Epitope Conservancy Analysis (ECA) tool was used to assess the similarity between the identified peptides and query sequences, based on defined parameters (Bui et al., 2007; Calis et al., 2013). For CD4+ T-cell peptides, those with IC50 values ≤ 100 (categorized as high, moderate, and low affinity) were identified using the MHC class-II binding prediction tool (Wang et al., 2010). The jobs on the said servers were run between 15 – 25 Feb 2023.

### Molecular docking, MD simulation, and visualization

Peptide docking was performed on GalaxyWeb (GalaxyPepDock) / ClusPro in peptide mode (Porter et al., 2017). Protein crystal structure (PDB ID: 7CW2) and predicted peptide, for T-Cell and B-Cell epitope prediction, were used for affinity prediction assessment. MD simulation was performed on GROMACS. For MD simulation, the models with docked peptides were visualized using Discovery Studio (v25.1.0.24284) from BIOVIA Dassault Systems, and PyMol (v3.0.3, Schrödinger LLC.).

#### Preparation of ligand peptides

A Previous literature search yielded ligand (here peptides) that scientists used for their docking analysis. The electrostatic potential in this context required accurate visualization through the usage of the PyMOL tool, which performed the tasks of normalizing geometry while adding hydrogen atoms and allocating charges and eliminating redundant atomic elements, if required.

#### Protein Preparation and energy minimization

The protein structures used for all three receptors originated from the Protein Data Bank. The AutoDock tool enabled the process for atom removal from the protein structure and the assignment of protein atom charges while adding missing atoms and carrying out energy minimization for structure optimization.

#### Molecular Docking Study

The protein-peptide docking was performed on GalaxyWeb (GalaxyPepDock) and ClusPro in peptide mode (Porter et al., 2017). PDB models were visualized either in PyMol or Discovery Studio.

The docking affinity in terms of bonding pattern analysis helped identify which model represents the best scenario while evaluating the candidate potential of peptides. The best predicted model was displayed to understand the mechanism of the process.

### Molecular Dynamics

GROMACS (GROningen MAchine for Chemical Simulations) performed molecular dynamics simulations for analyzing the dynamic movements of the target protein structure. The simulation relied on the force field AMBER99SB-ILDN to parameterize the system before placing the protein in a cubic water chamber using TIP3P liquid while leaving a 1.0 nm buffer between the protein and box walls. The system required appropriate counterions for its neutralization as well as the simulation of physiological conditions. A systematic process of energy minimization resolved steric issues while enhancing system geometrical performance. The simulation passed through two stages for system equilibrium: NVT phase and NPT phase, which maintained constant number, volume, and temperature conditions before the production run. The production MD simulation took place at 300 K temperature and 1 bar pressure while applying a combination of Berendsen thermostat and Parrinello-Rahman barostat. The simulation utilized a 2-femtosecond time step under Periodic Boundary Conditions with Particle Mesh Ewald method for computing electrostatics at long distances. Structural stability together with flexibility measurements (RMSF) and conformational alterations served as the analysis criteria for the trajectories recorded during simulation time.

### Gene designing

The selected full-length sequence of the E2 gene from the S27 African strain of the Chikungunya virus was obtained from NCBI (GeneBank Accession No. AF339485.1; Supplementary Table 1). The E2 gene sequence of the Chikungunya virus (1287bp) was chemically synthesized by GenScript (New Jersey, USA). The synthesized sequence was inserted after a BamHI sequence at the 5’ end, followed by the start codon, E2 gene and a _6x_His tag before a stop codon and a Hind III sequence at the 3’ end. In the entire study, three vectors were used for cloning and expression, viz., pUC57, pET28a(+) and pCAMBIA1302.

### Expression of rE2 in E. coli Cloning of rE2

The gene cassette was inserted into the cloning vector pUC57 using BamHI and HindIII restriction sites, and the resulting construct was then grown. Subsequently, the plasmid was isolated, and the rE2 gene was amplified using Taq polymerase (NEB, England) with specific forward (5’ GGATCCATGAGCACCAAGGACAACT 3’) and reverse primers (5’ AAGCTTTTACGCTTTAGCTGTTCTG 3’). The resulting PCR product was analyzed on a 1% agarose gel. The rE2 was then subcloned into pET28a(+) expression (under T7 promoter) vectors. The constructs were transferred into the BL21(DE3)pLysS strain of *E. coli*. A successful cloning was confirmed by restriction digestion.

#### Expression of rE2 protein

For the expression of rE2 protein using pET28a(+), transformed *E. coli* were grown overnight at 37[ using 50 mg/mL kanamycin. This overnight culture was then inoculated into 50 mL LB medium and incubated at 37°C under shaking conditions until the bacterial culture reached an optical density (OD) of 0.4-0.6. Expression of rE2 was induced by iso-propyl thio-beta-D-glucosidase (IPTG) at a concentration of 1 mM for 4 h. Single colonies were sub-cultured and screened to check the rE2 gene. Before the induction and 16 h (overnight) of induction, samples were processed for protein extraction. Protein was checked on the SDS-PAGE to locate the newly expressed protein band.

The culture was pelleted in a microtube and suspended in 1XPBS, followed by a brief sonication. Tubes were spun at 10,000 rpm for 15 min at 4°C. Both supernatant and pellet were run on the SDS-PAGE for verification. The sonicated supernatant was further processed for Ni-NTA affinity chromatography.

#### Isolation of protein from E. coli

Total protein from *E. coli* was isolated, and rE2 was isolated on the Ni-NTA column with the help of His-tag using the method mentioned by the provider (Handbook - QIAexpressionist^TM^, Qiagen). The cell pellet obtained upon induction was lysed in a Lysis solution supplemented with 1× Protease Inhibitor Cocktail and comprising 25 mM Tris pH 7.3, 8 M urea, 5 mM β-mercaptoethanol, and 300 mM NaCl. After centrifuging the lysate for 30 minutes at 4°C at 12,000 rpm, the supernatant containing the protein was put onto a column with 1 mL of Ni-NTA resin, and it was left to bind for 15 minutes at room temperature. Wash buffers (lysis buffer containing 8 M urea supplemented with imidazole (20, 50, and 100 mM)) were used to wash the column. Finally, an elution solution supplemented with imidazole (100 mM imidazole, 250 mM imidazole, 500 mM imidazole, 750 mM imidazole, and 1M imidazole) and lysis buffer with 8 M urea was used to wash the column. To estimate purity, the washed and eluted fractions were subjected to analysis on 12% SDS-PAGE.

### Expression of rE2 protein in *Nicotiana tabacum* plants

#### Subcloning of rE2 gene into pCAMBIA1302

The expression of Chikungunya virus rE2 protein in *Nicotiana tabacum* plants involved subcloning of the rE2 gene into the MCS of pCAMBIA1302 vector (under the control of CaMV promoter) using BamHI and HindIII.

#### *A. tumefaciens* transformation and plant infiltration

The recombinant vector pCAMBIA1302-rE2 was mobilized into *Agrobacterium tumefaciens* (EHA105) using the liquid N_2_ and heat shock method. Competent cells were prepared using the CaCl_2_ method, and confirmation of transformation was achieved through colony PCR.

Seeds of *Nicotiana tabacum* were obtained from the Indian Agricultural Research Institute (IARI), Pusa, New Delhi, India. Seeds were surface sterilized and germinated on MS media. Approximately a hundred healthy, physically uniform tobacco seeds were washed with Tween 20 detergent, surface sterilized with 70% ethanol for 45 seconds each, and further disinfected with NaClO (1%) and mercuric chloride (0.1%) for 45 seconds. After extensive washing with double-distilled water, the seeds were dried on sterile filter paper and plated onto jam bottles containing MS media. Germination occurred under controlled conditions with a photoperiod and temperature of 16 h at 28[/8 h 24[ (day/night). Once reaching the four-leaf stage, the plants were transferred to the greenhouse and grown as mentioned by Malhotra et al. (2016). Agroinfiltration in *Nicotiana tabacum* followed the protocol of Aguilar et al. (2016).

*A. tumefaciens* EHA105 strain, cultured in YEM media with 50 mg/mL kanamycin and 30 mg/mL rifampicin, was used. Single colonies containing pCAMBIA1302 vector–rE2 were inoculated into YEM media and grown at 28°C with shaking at 200 rpm. An aliquot of 1.0 mL was inoculated into 10 mL YEM media and grown overnight to reach an OD_600_ of 0.6. Bacteria were spun at 15,000 rpm and then resuspended in an induction buffer containing MMA (10 mM MgCl_2_, 10 mM MES (2-morpholinoethanesulfonic acid, and 200 μM acetosyringone). The mixture was gently swirled until a clear homogenous culture formed, and incubation was done for 2 to 4 h at 24°C with gentle shaking. Infiltration was done into the eight-week-old plants with 1 mL syringes (without a needle) to infiltrate the mixture via the abaxial surface of the leaf. Two to four fully expanded leaves per plant were used for infiltration.

### Fluorescence monitoring in plants

Fluorescence monitoring in plants involved collecting samples at 3dpi and 8dpi. Fluorescence was monitored in wild-type, 3^rd^ dpi, and 8^th^ dpi samples using a Leica confocal TCS SP5 microscope. Parameters included light: LED epi, exposure time: 3-120 sec, color: Blue, and filter name: 530 bp.

### Real-time expression studies in tobacco leaf using qPCR

RNA isolation was done by the TRIzol method as detailed in Sangha et al. (2010). Samples were collected from wild-type plants at 3rd and 8th days post-infection (dpi). Synthesis of cDNA was done using the Verso cDNA Kit. The concentration of cDNA was determined using a nanodrop. Primers for RT-PCR were designed using IDT primer quest tools. The E2 gene, a 102-base pair fragment (amplicon), was amplified using the forward primer (5’ CCATAGTCCCGTAGCACTAGAA3’) and reverse primer (5’ TGGCTATCATCCGTCCCTATT 3’). Its expression was quantified relative to actin protein, with control forward primer (5’TTGACTATGAGCAGGAGC3’) and reverse primer (5’CAGCTTCCATTCCGATCA3’).

Expression levels were determined by the cycle threshold (Cq) and analyzed using the ΔΔCt method. ΔCt was calculated as (transgene – actin), and ΔΔCt was calculated as (ΔΔCt of transgenic - ΔΔCt of actin). Standard error was computed, and a graph depicting the expression level of transiently transformed samples, which were imported into Microsoft Excel, was drawn.

### Protein extraction from tobacco leaf

*N. tabacum* leaf samples, collected on the 3rd day post-Agro-infiltration, were snap-frozen in liquid nitrogen and immediately crushed in a mortar pestle (ice cold) in 2.0 mL chilled PBS. Total protein extraction from 100 mg of transiently transformed tobacco leaf utilized 1X extraction buffer (0.1% Triton-X, 150 mM NaCl, 100 mM sodium phosphate at pH 6.8, and protease inhibitors). Protein content was estimated using the Bradford method, with BSA protein as the standard. A 200 µL of SDS-PAGE gel loading dye per mL of extract was added, followed by boiling for 3 min and quick chilling.

### Isolation of rE2 from tobacco

The protein was purified using the previously mentioned protocol employing Ni-NTA affinity chromatography. Subsequently, the rE2 protein underwent dialysis against phosphate-buffered saline (PBS) before downstream testing. The recombinant purified rE2 protein was utilized for further studies.

### In vivo studies

#### Immunogenicity studies in BALB/c mice

All animal studies were conducted at Jamia Hamdard, New Delhi, following proper set guidelines. The studies complied with Animal Welfare Act regulations and were approved by the Institutional Animal Ethics Committee, proposal no. 2088, adhering to the guidelines of CCSEA Ministry of Social Justice, Govt. of India, which is adopted by the Institutional Animal Ethics Committee at Jamia Hamdard, New Delhi, India. Two independent animal immunization experiments were conducted using BALB/c mice aged 6–8 weeks. One experiment utilized purified recombinant protein from E. coli, and the other used recombinant E2 protein of CHIKV from *Nicotiana tabacum*.

Four different formulations with the recombinant rE2 were used as follows:

– Group A: Normal Saline
– Group B: Recombinant rE2
– Group C: Recombinant rE2 (Complete Freund adjuvant + Incomplete Freund adjuvant)
– Group D: Recombinant rE2 + Alum adjuvant

In Group A (control group), BALB/c mice were injected with a sterile normal saline solution. In Group B, mice were immunized with purified recombinant E2 protein. In Group C, a primary injection with 100 µg of antigen in Complete Freund’s adjuvant (CFA) was administered subcutaneously on Day 1. The 1st booster at Day 21 involved 50 µg of antigen in Incomplete Freund’s adjuvant (IFA), the 2nd booster at Day 42 included antigen in IFA, followed by serum collection. The 3rd booster at Day 62 consisted of 50 µg of IFA.

In Group D, CHIKV-E2 protein mixed with alum adjuvant (Thermo Fisher Scientific, USA) was administered, and booster doses were given similarly as with complete Freund adjuvant and incomplete Freund adjuvant. After allowing the separated blood samples to coagulate for two to three hours at room temperature, the serum was separated using a centrifuge.

### Titre Estimation E2-Specific ELISA

Detection of antigen-specific IgG antibodies was conducted using ELISA. For coating, 1.0 μg/mL of protein was prepared in carbonate coating buffer and distributed into the ELISA plate in a 100 μL volume. The plate (Sigma, United States) was incubated overnight at 2-8[. After the overnight incubation, the plate was washed three times with 300 μL of PBST (PBS with 0.1% Tween 20). About 200 μL of 1% BSA prepared in PBS was added to the plate and incubated for 1 hour at room temperature. The plates were washed three times with 300 μL of PBST.

Different dilutions of mouse sera were prepared in 0.1% BSA-PBS. About 100 μL was added to the respective wells and incubated for 1 h at room temperature. The plates were washed three times with 300 μL of PBST. About 100 μL of 1:10000 diluted anti-mouse IgG-HRP conjugate was added to the plate and incubated for 1 h at room temperature. The plates were washed three times with 300 μL of PBST. About 100 μL of TMB substrate was added to the plate and incubated for 20 min at room temperature. Then, 50 μL of stop solution was added to the plate to stop the reaction. Readings were taken at 450 nm with 570 nm using a microplate reader. Dilutions showing above 0.3 OD were considered titers. If the OD value of the tested sample exceeded the OD of the negative control by a factor of 3, the samples were considered positive.

### Western blot analysis

To assess the antigen-antibody specificity of the rE2 protein isolated from *E. coli* and tobacco (3rd day post-Agro-infiltration), a western blot analysis was performed. First, SDS-PAGE was run at 50V for the initial 30 min, followed by 100V for approximately 2 h, and then transferred onto a nitrocellulose membrane. The membrane was blocked in TBS containing BSA (5%) overnight at 40 [. Subsequently, antisera (anti-rE2 mice IgG) at a dilution of 1:1000 was incubated for 2 hours at a temperature of 37[. The membrane was then washed with TBS containing Tween 20 (0.05%) and further incubated with horseradish peroxidase-conjugated anti-mouse IgG antibodies at a dilution of 1:5000 for one hour at 37[. Following this, the blotting membrane was washed with TBS-T buffer and stained with a solution of 3,3’,5,5’-tetramethyl-benzidine containing H_2_O_2_ until a colored solution appeared. Water was added to terminate the reaction immediately. The gel pattern was then digitally documented on ChemiDoc (Bio-Rad, USA).

## RESULTS

This study aimed to clone and express the recombinant CHIKV E2 gene to produce recombinant E2 protein both in *E. coli* and a plant system (*N. tabacum*) and test immunogenicity in the BALB/c mice. Before cloning and expression of CHIKV E2, comprehensive computational studies were performed using the crystal structure from PDB. The objective was significantly achieved, and the results are presented in this section.

### The study made a significant contribution to the structural features of CHIKV E2

Amino acid sequence of the E2 protein of chikungunya virus (S27-African strain, UniProt ID Q8JUX5) is presented in the supplementary Table 2. The amino acid frequency chart (Figure 1) shows the frequencies of each amino acid residue in the E2 protein. Threonine (T) was most abundant (38 residues), whereas tryptophan (W) was least abundant (5 residues).

**Figure 1:**
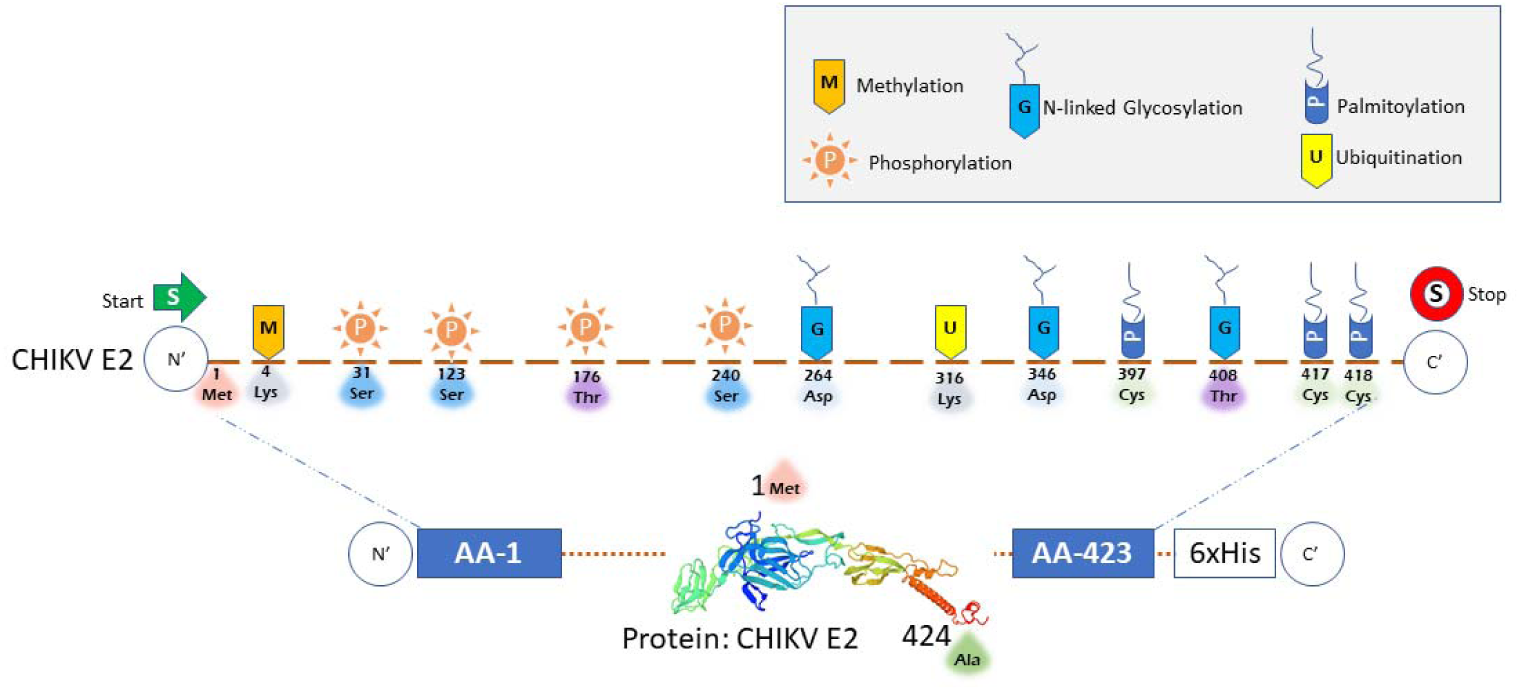
Possible post-translational modification (PTM) sites in the CHIKV E2 protein as predicted by MuSiteDeep Portal. Computational assessment was done based on predicted PTM sites in the human proteome using the PTMGPT2 portal. In the bacterial and plant systems where recombinant E2 was expressed, many residues might be left PTM-vacant and could be humanized after being injected into mammalian (human/mouse) systems. Details of such sites are available in Table 1.

**Table 1:**
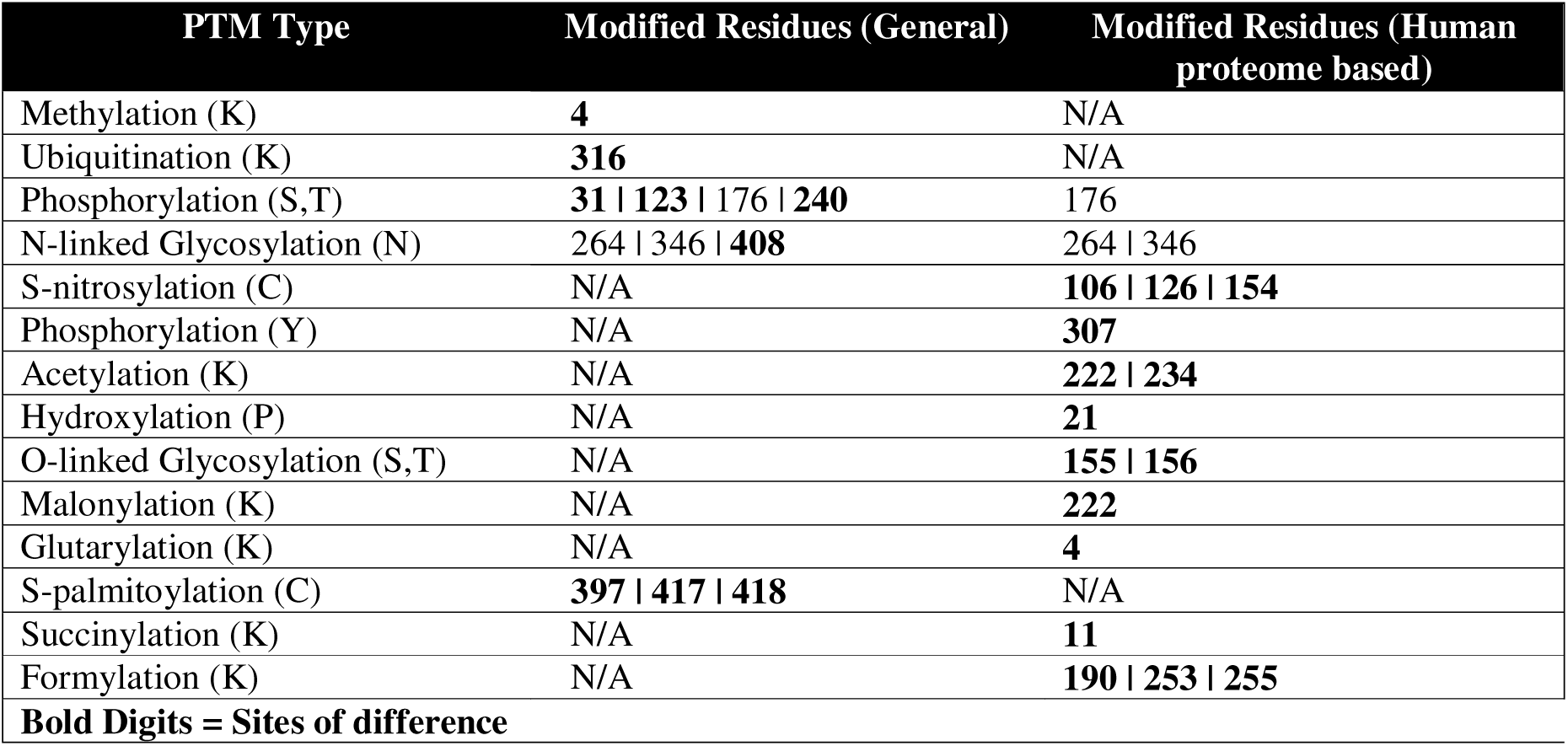
Comparative PTM prediction summary of modified residues that might take place in bacteria and plants versus.

Based on the amino acid frequency chart (Supplementary Figure 1), the E2 protein had a distinct composition. The most abundant amino acids are Threonine, Valine, Proline, Leucine, and Glycine, suggesting a structure rich in hydrophobic regions, specific turns, and flexible areas. The high frequency of Threonine and Valine, in particular, points to a significant presence of non-polar or hydrophobic residues that likely contribute to the protein’s core or its interactions with other non-polar environments, such as a cell membrane. Additionally, the elevated counts of Proline and Glycine hint at a complex and potentially dynamic structure, with Proline introducing kinks and Glycine providing flexibility, both essential for forming specific loops and turns.

#### Phylogenetic assessment

Phylogenetic analysis provided a comprehensive understanding of related protein sequences affected by genetic changes. MEGA12’s phylogenetic analysis program constructed the tree using a neighbor-joining strategy, showing that the E2 protein of the Chikungunya virus has retained distinctive domains, and there is minimal variation in the length of amino acids across different Chikungunya virus serotypes. The closest serotypes to the E2 protein considered in the present study were extremely close to the serotypes found in the majority of Chikungunya fever-affected regions, including India, Senegal, Tanzania, Japan, Italy, Sri Lanka, West Africa, etc. (Supplementary Figure 2).

#### Characteristics of CHIKV E2 3D model

Individual subunit of viral capsid, E2, was re-modeled on the portal SWISSMODEL, taking 6NK5 (PDB ID) as a template. Multicrit analysis on Molprobity Version 4.4 (SwissModel Portal) showed a MolProbity score of 1.50, a clash score of 0.31, 88.49% and 2.88% per cent of Ramachandran favored and outliers, respectively. There were 2.47% rotamer outliers, 21 C-beta deviations, zero bad bonds, 44 out of 4584 (0.96%) bad angles, and 3 out of 387 (0.775%) twisted non-proline. The model (Supplementary Figure 3), the Ramachandran plot (Supplementary Figure 4), and the multicrit table (Supplementary Table 3) describe the stability and possible dynamics of protein structure.

### Comprehensive PTM features of CHIKV E2

A comprehensive analysis revealed interesting facts about the CHIKV E2 protein (Figure 1). VPTMdb, a viral PTM database, showed that the top ten E2 sequences from various serotypes had a similar pattern of PTMs to Q8JUX5, which was taken as the model protein sequence. In the E2, based on computational probability and human proteome data, there was one methylation site (Lys4), four phosphorylation sites (Ser31, 123, 240 and Thr176), three N-linked glycosylation sites (Asp264 and 346, and Thr408), one ubiquitination site (Lys316), and three palmitoylation site (Cys397, 417 an 418) were predicted (Figure 1) using MuSiteDeep Portal. Compared to the human PTMs (human proteome algorithm; PTMGPT2), there was additional methylation (Lys4), phosphorylation (Ser31, 123, and 240), N-linked glycosylation (Thr408), and three palmitoylation site (Cys397, 417 an 418) were predicted. Upon arrival in the human, three S-nitrosylation (Cys106, 126 and 154), one phosphorylation (Tyr307), two acetylation (Lys222 and 234), one hydroxylation (Pro21), O-linked glycosylation (Ser155 and Thr156), malonylation (Lys222), glutarylation (Lys4), succinylation (Lys11), and formylation (Lys153, 253 and 255) was predicted in individual or cross-talk manner.

Comparative PTM prediction summary of modified residues, which might take place in bacteria and plants, versus mammal (human/mouse) reflects that there is a lot of opportunity for recombinant E2 (expressed in *E. coli* and *N. tabacum*) to get humanized PTMs, as the E2 of the actual Chikungunya virus might undergo in the human body (Table 1).

### CHIKV E2 epitope prediction

There are several reports on the prediction of B-cell and T-cell epitopes. However, with emerging tools and improved versions of epitope predictions, there were slight or significant differences have been observed. Furthermore, a comprehensive docking analysis for each peptide with MHCs (both Class I and Class II), with novel visualization, makes this study unique.

#### B-cell epitope prediction

The B-cell epitopes comprise peptides that can easily be used to take the place of antigens for immunizations and antibody production (Supplementary Figure 5 A-E). We found B-cell linear epitopes in the surface glycoprotein of the S27-African strain of Chikungunya virus, which may be capable of inducing the desired immune response as B-cell epitopes. In total, there were 11 peptides with lengths varying from 6 to 19 residues as predicted by Emini Surface Accessibility Prediction (IEDB) (Supplementary Figure 5A and Supplementary Table 4), 13 peptides varying from 6 to 24 residues were predicted by Bepipred Linear Epitope Prediction 2.0 (Supplementary Figure 5B and Supplementary Table 5), five peptides were predicted by Chou & Fasman beta turn prediction tool (Supplementary Figure 5C and Supplementary Table 6), peptides predicted on parameters of hydrophilicity, flexibility, accessibility, and turns by Parker Hydrophilicity Tool are mentioned in Supplementary Figure 5D and Supplementary Table 6, and 16 peptides varying from 6 to 17 residues were predicted by Kolaskar & Tongaonkar Antigenicity tool (Supplementary Figure 5E and Supplementary Table 7).

#### T-cell epitopes prediction for CHIKV E2

T-cell epitope prediction for the structural E2 protein of the Chikungunya virus using TepiTool resulted in the identification of peptides from MHC Class-I alleles with IC50 values ≤ 100 nm, showing high binding affinity. The total score of each peptide–HLA interaction was considered, with a higher score indicating greater processing efficiency. The peptides HPHEIILYY (P1), RSMGEEPNY (P2), KNQVIMLLY (P3), STKDNFNVY (P4), and VTWGNNEPY (P5) were found to bind with various MHC Class-I molecules (Supplementary Table 8). Additionally, epitope conservancy analysis showed maximum identity (100%) for these peptides.

For MHC class II potential peptide epitopes, three CD4+ T-cell peptides were predicted, and four peptides with high binding affinity (IC50 ≤ 50) were selected. These peptides (DNFNVYKATRPYLAH, RAGLFVRTSAPCTIT, and YYYELYPTMTVVVVS) interacted with various HLA-DRB-1 molecules, as detailed in the Supplementary Table 9.

#### Molecular docking of selective epitopes

As predicted, 5 epitopes were docked with CD8+ (MHC Class I; Figure 2A-E) and three with CD4+ (MHC Class II; Figure 3A-C). The docking (peptide-protein) showed impressive models of affinity.

**Figure 2.**
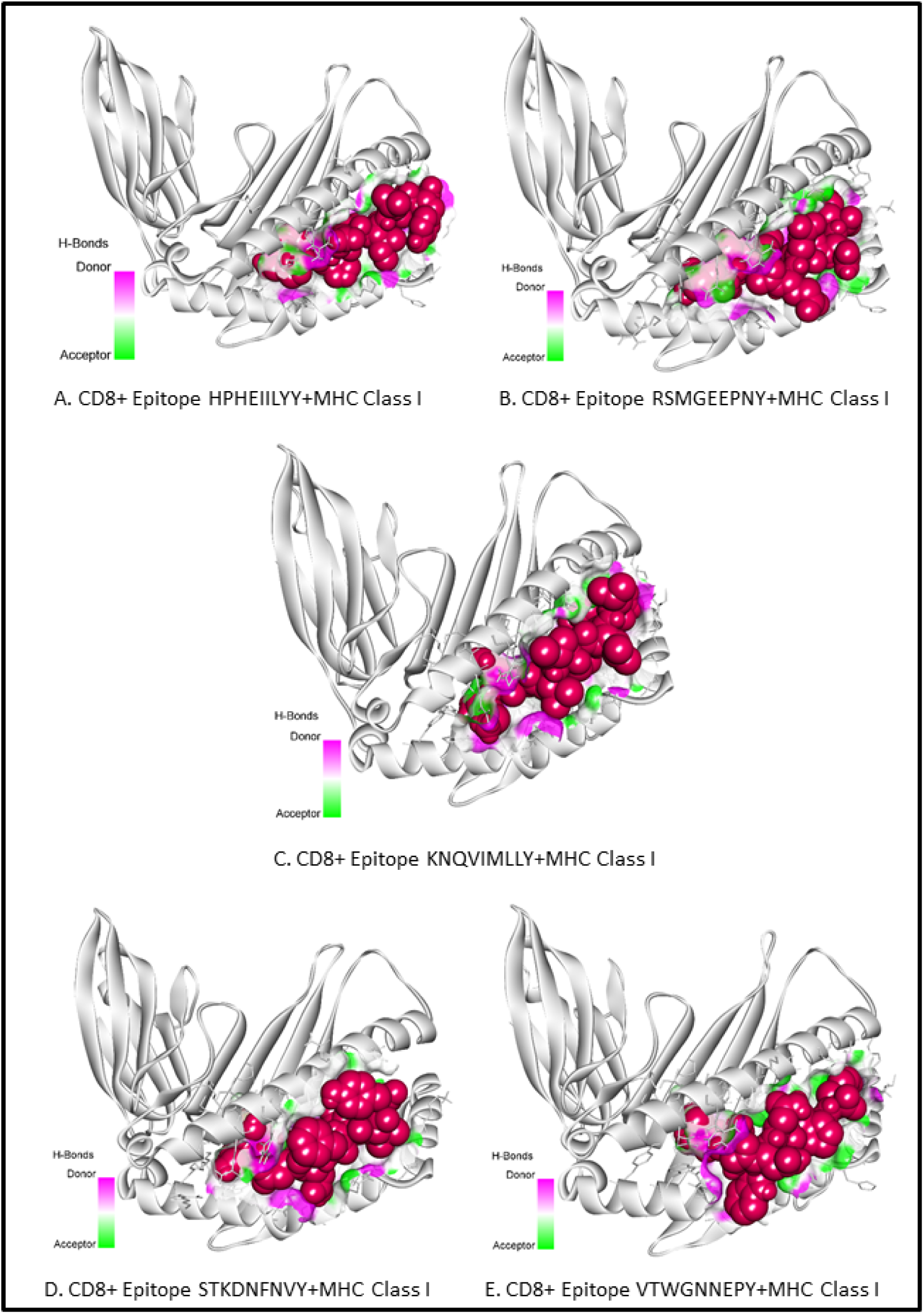
**(A-E):** Display of molecular peptide to protein docking of different epitopes with CD8+ MHC (Class I). Red balls show epitope residues. Details of all interactions are provided in Supplementary Tables (10-14).

**Figure 3.**
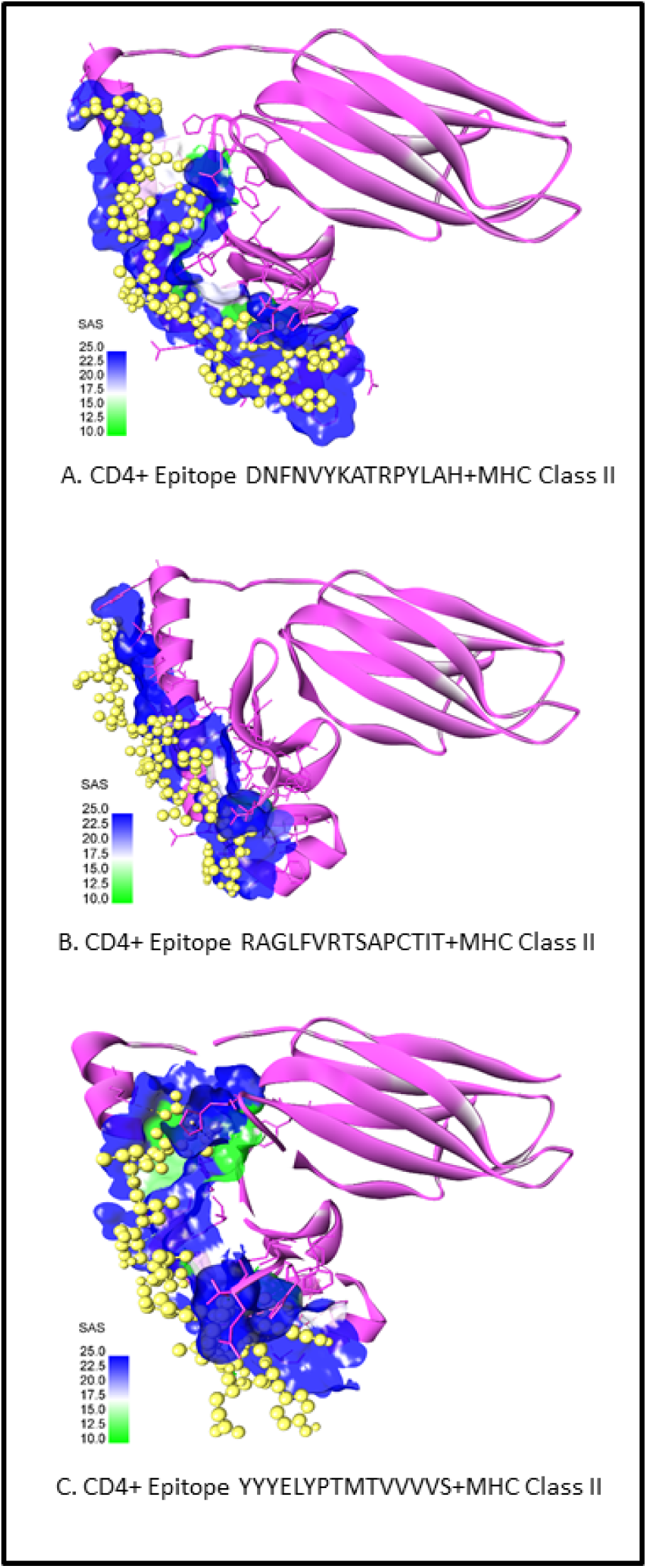
**(A-C):** Display of molecular peptide to protein docking of different epitopes with CD4+ MHC Class II. Yellow discs show epitope residues. Details of all interactions are provided in Supplementary Tables (15-17).

Interaction of predicted CD8+ T-cell epitope “HPHEIILYY” (Supplementary Table 10), “RSMGEEPNY” (Supplementary Table 11), KNQVIMLLY (Supplementary Table 12), STKDNFNVY (Supplementary Table 13) and VTWGNNEPY (Supplementary Table 14) in CHIKV E2 with MHC class-I alleles with varied affinity of IC50 showed enormous types of interactions.

On the other hand, Interaction of predicted CD4+ T-cell epitope DNFNVYKATRPYLAH (Supplementary Table 15), RAGLFVRTSAPCTIT (Supplementary Table 16), and YYYELYPTMTVVVVS (Supplementary Table 17) in CHIKV E2 with MHC class-II alleles having affinity IC50 < 100 were found to have good affinity but lesser diversity of interactions compared to MHC class I.

### Molecular dynamics (MD) simulation studies

MD simulations offer a detailed and dynamic view of peptide-protein interaction at the atomic level. Thus, it provides rigorous data on affinity calculations based on pose refinement. We performed MD simulations to investigate the structural stability and dynamic behavior of the human leukocyte antigen HLA-DRB1 complexed with the peptide YYYELYPTMTVVVVS over a 100-ns trajectory. Peptide 3 (YYYELYPTMTVVVVS) was the best candidate for simulation with HLA-DRB1*13:02 due to the lowest IC50 (64.52). The choice of the “best” peptide for MD simulation was allele-specific; we selected the peptide-allele pair with the lowest IC50 value for their particular study.

The initial conformation of the HLA-DRB1-peptide complex is shown in Figure 3 (C). Further, a three-dimensional representation of the HLA-DRB1 (Figure 4A-E) protein (density blue colored) bound to the peptide YYYELYPTMTVVVVS (displayed in yellow color) (Figure 4A). The starting conformation of the molecular dynamics simulation, illustrating the peptide nestled within the binding groove of the HLA-DRB1 molecule, has been demonstrated (Figure 4B-E).

**Figure 4.**
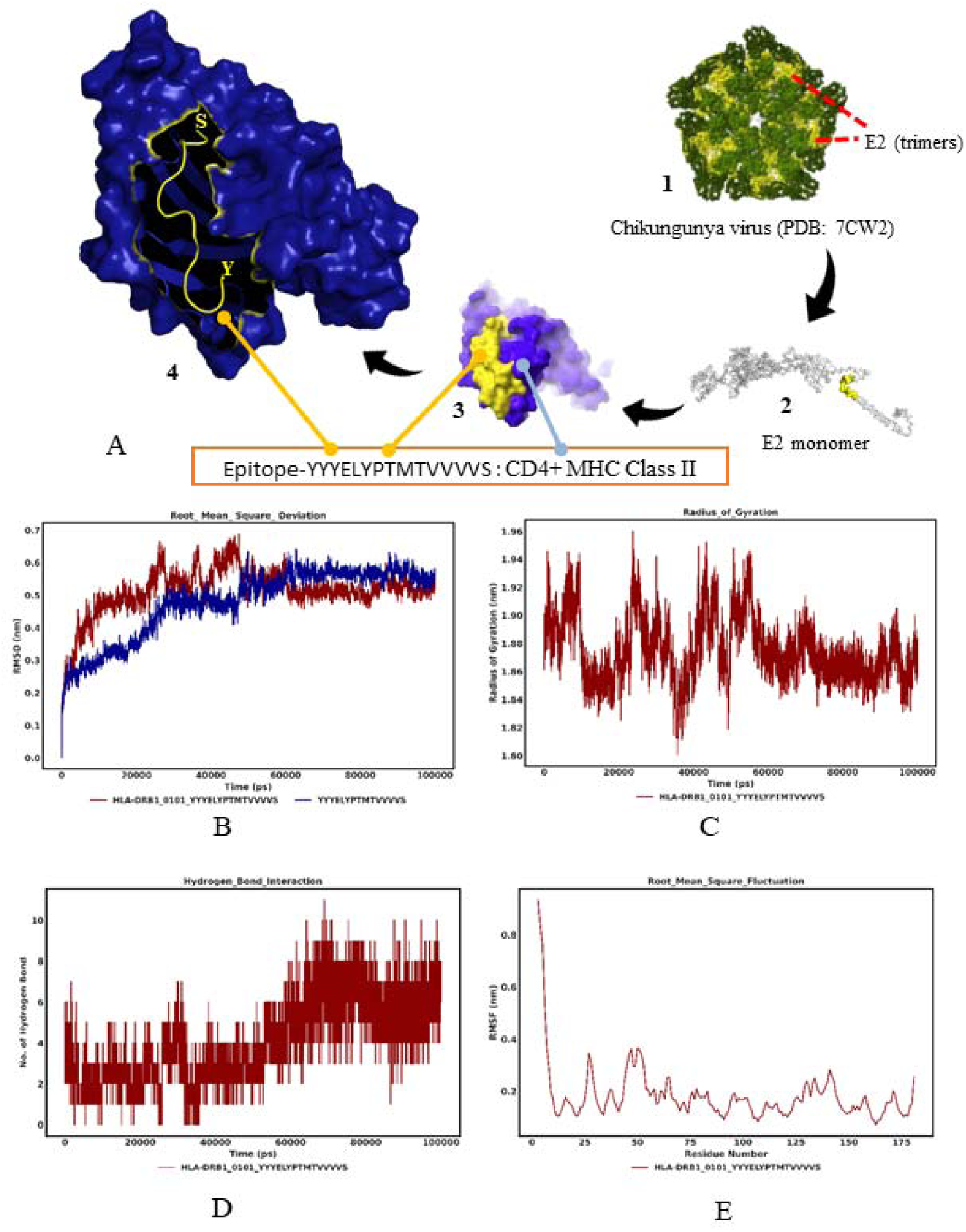
**(A-E): A.** Crystal structure of chikungunya virus (PDB ID: 7CW2); presented in full model (A.1), isolated structural protein E2 (A.2), a predicted epitope in surface-to-surface association with one of the antigen-presenting HLA-DRB1 (A.3), and a zoom-in model of i*n silico*-docked between HLA-DRB1 protein (A.4) (density blue colored surface) and the peptide YYYELYPTMTVVVVS (displayed in yellow color). **B.** Root Mean Square Deviation (RMSD) of the HLA-DRB1-Peptide Complex and Peptide over 100 ns Molecular Dynamics Simulation, indicating system equilibration and structural stability. **C.** Radius of Gyration (Rg) of the HLA-DRB1-Peptide Complex over 100 ns Molecular Dynamics Simulation. Fluctuations in Rg indicate changes in the overall shape and density, while a stable plateau suggests that the complex maintains a consistent globular structure. **D.** Intermolecular Hydrogen Bond Interactions between HLA-DRB1 and Peptide YYYELYPTMTVVVVS over 100 ns Molecular Dynamics Simulation. This plot shows the total number of hydrogen bonds formed between the HLA-DRB1 protein and the peptide YYYELYPTMTVVVVS as a function of simulation time (in picoseconds, ps). The dynamic nature of hydrogen bond formation and breakage is reflected in the fluctuations observed over the trajectory. **E.** Root Mean Square Fluctuation (RMSF) of HLA-DRB1-Peptide Complex Residues plotted against the residue index. Some residues exhibit distinct flexibilities.

### Overall System Stability: Root Mean Square Deviation (RMSD) Analysis

The stability of the HLA-DRB1-peptide complex throughout the 100 ns simulation was assessed by calculating the Root Mean Square Deviation (RMSD) of the protein backbone atoms relative to the initial structure (Figure 4B). The RMSD of the entire HLA-DRB1-peptide complex (red line) showed an initial increase during the first ∼10 ns, reaching approximately 0.45 nm, indicative of an equilibration phase. Subsequently, the RMSD gradually increased, fluctuating between 0.5 nm and 0.65 nm up to approximately 50 ns, before stabilizing and fluctuating around an average of 0.55 nm for the remainder of the simulation (50-100 ns). This stable plateau suggests that the overall complex achieved a well-equilibrated and stable conformation.

Concomitantly, the RMSD of the peptide YYYELYPTMTVVVVS alone (blue line), when aligned to its initial position, closely mirrored the trajectory of the entire complex. The peptide’s RMSD also exhibited an initial rise followed by a more gradual increase, reaching values around 0.5 nm by 50 ns and subsequently plateauing alongside the complex’s RMSD. The synchronized behavior of the peptide and complex RMSD indicates that the peptide remained stably bound within the HLA-DRB1 binding groove and did not undergo significant independent conformational excursions or dissociation from the protein during the simulation.

### Compactness Assessment: Radius of Gyration (Rg)

The global compactness of the HLA-DRB1-peptide complex was monitored through the Radius of Gyration (Rg) over the 100 ns simulation (Figure 4C). The Rg values for the complex fluctuated between approximately 1.80 nm and 1.96 nm throughout the entire trajectory. No sustained upward or downward trend was observed, indicating that the HLA-DRB1-peptide complex maintained its overall folded and compact structure. The observed fluctuations are consistent with inherent thermal motions and dynamic breathing of a protein-peptide complex in solution.

### Interfacial Interactions: Hydrogen Bond Analysis

The number of hydrogen bonds formed between the HLA-DRB1 protein and the peptide YYYELYPTMTVVVVS was calculated over the simulation time to quantify the stability of the intermolecular interaction (Figure 4D). The hydrogen bond count displayed dynamic fluctuations, ranging from 0 to over 10 bonds throughout the simulation. In the initial phase (∼0-50 ns), the number of hydrogen bonds was highly variable, sometimes dropping to lower counts but frequently reaching 5-7 bonds. Notably, from approximately 50 ns onwards, the hydrogen bond network appeared more robust, with the number of bonds often fluctuating between 8 and 10, suggesting a stronger and more consistent interaction in the latter half of the simulation. This sustained presence of multiple hydrogen bonds emphasizes the stable anchoring of the peptide within the HLA-DRB1 binding site.

### Per-Residue Flexibility: Root Mean Square Fluctuation (RMSF) Analysis

To identify flexible and rigid regions within the HLA-DRB1 protein when bound to the peptide, the Root Mean Square Fluctuation (RMSF) was calculated for each residue (Figure 4E). The RMSF profile revealed varying degrees of flexibility across the protein sequence. Regions exhibiting lower RMSF values indicate relatively rigid structural elements, likely corresponding to stable secondary structures comprising the peptide-binding groove. On the other hand, higher RMSF values were observed in regions around many residues, suggesting greater conformational flexibility, often characteristic of loop regions or termini exposed to the solvent. These findings provide insights into the intrinsic dynamics of the HLA-DRB1 protein in complex with the peptide.

### Cloning and expression of CHIKV E2

The nucleotide sequence of the S27-African strain of the CHIKV E2 gene (GeneBank Accession No. AF339485.1; Supplementary Table 1) was optimized and chemically synthesized by GenScript (New Jersey, USA) and cloned into different vectors to express in *E. coli* and *N. tabacum*.

#### Cloning in E. coli

The recombinant E2 (rE2) gene was successfully cloned into a pUC57 cloning vector in the BamHI/HindIII restriction sites (Supplementary Figure 6). The PCR product was run on a 1% agarose gel, which confirmed successful gene amplification (Figure 5A). Further, the cloning in pUC57 was confirmed by restriction digestion using the same set of restriction enzymes BamHI and HindIII (Figure 5B) on the agarose gel. The rE2 gene was then subcloned into the pET28a(+) expression vector (Supplementary Figure 7). The digested fragments were loaded onto a 1% agarose gel, revealing two distinct bands – one representing the uncut pET28a(+) vector backbone and the other indicating the released insert (Figure 5C).

**Figure 5.**
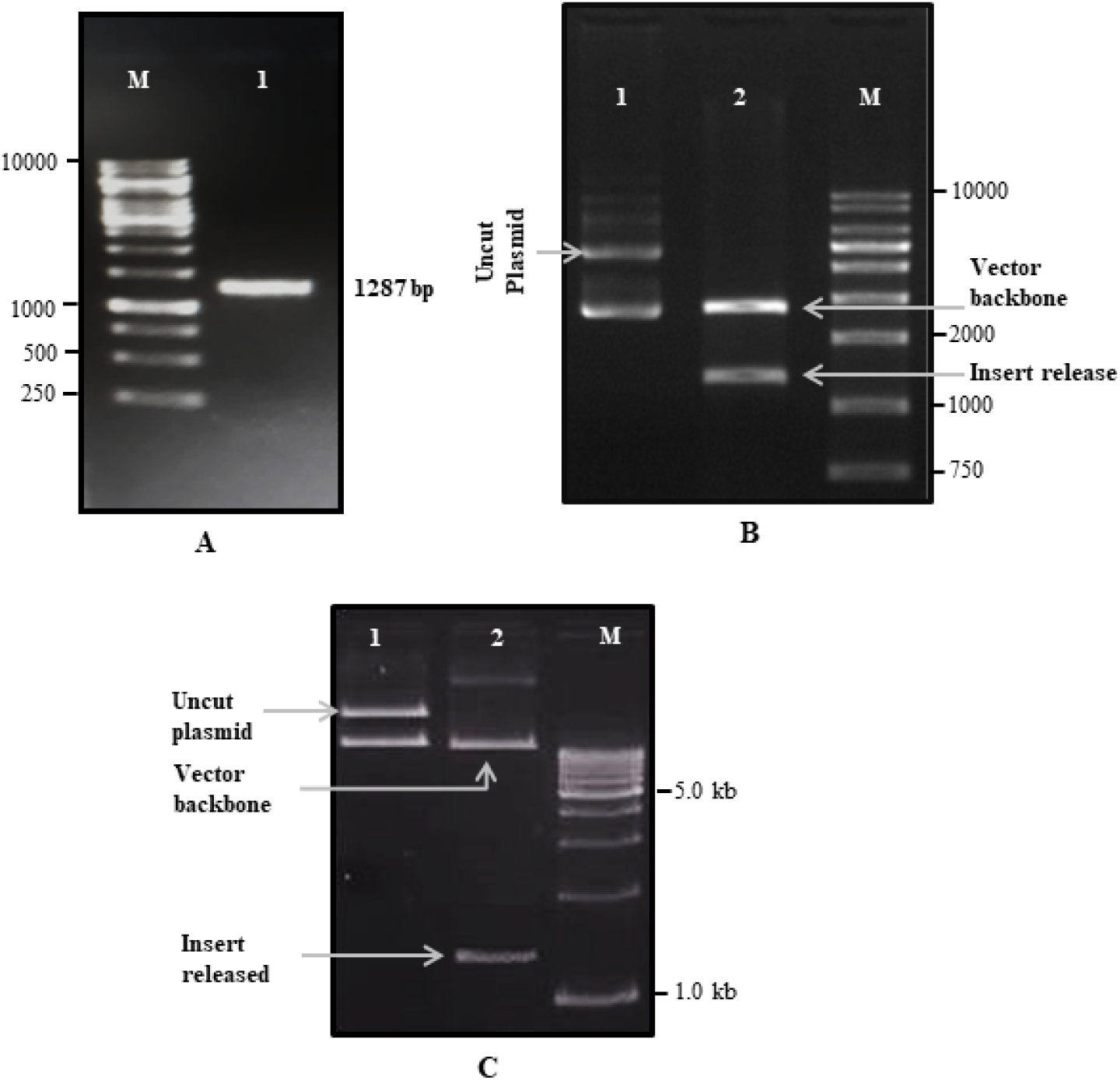
**(A-C): A.** PCR Amplification of rE2 gene of Chikungunya virus: Lane M (Marker); Lane 1 PCR-amplified rE2 gene of Chikungunya virus. The cloning in pUC57 was confirmed by restriction digestion using the same restriction enzymes BamHI and HindIII. **B.** Restriction digestion of the rE2 gene of Chikungunya virus from pUC57 shows confirmation of the gene on 1% agarose gel. Lane 1, uncut Plasmid; Lane 2, digested plasmid containing the vector backbone and insert release, and Lane M (Marker). **C.** Confirmation of rE2 cloning by restriction digestion; **Lane 1** represents the uncut plasmid pET28a(+) containing the E2 gene, **Lane 2** describes the double digestion released insert, and the vector backbone. **Lane M** represents Marker.

Positive clones were used for the transformation of the BL21DE3(plysS) strain of *E. coli* with the construct (Figure 6A). The confirmation of cloning and growth of BL21(DE3)pLysS strain was observed by growth on a Petri plate, where white-coloured colonies were obtained (Figure 6B).

**Figure 6.**
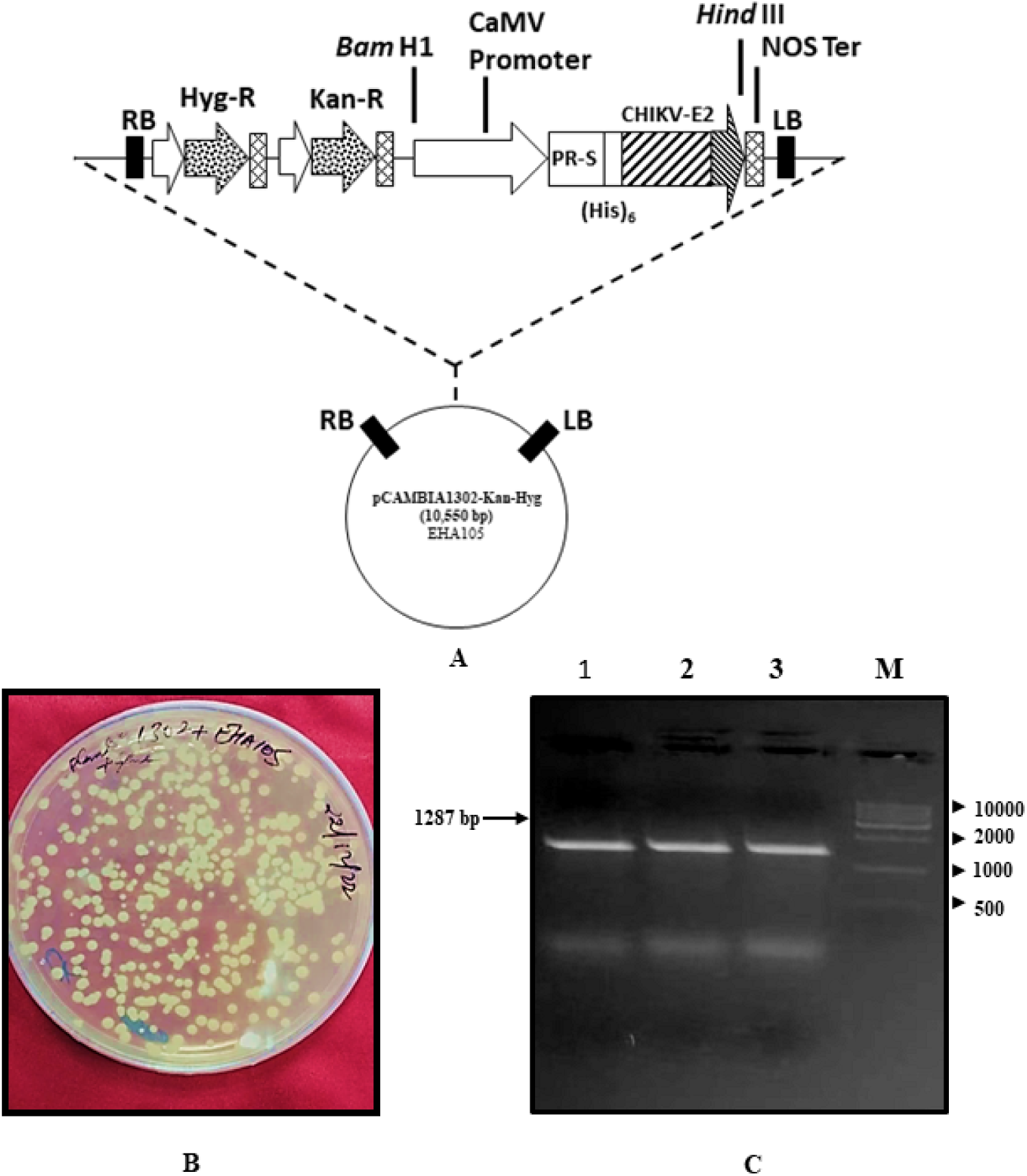
**(A-C): A.** Plant expression cassette used for expressing recombinant CHIKV-E2 (rE2) protein in the leaves of *N. tabacum* plant used in this study. The expression sequence for _6x_His tag was present just before the stop codon. **B.** Screening of transformants shown by the growth of EHA105 strain on YEM plates containing selection media with 50 mg/mL kanamycin and 30 mg/mL rifampicin. **C.** Confirmation of 1287 bp band on 1% agarose gel from Colony PCR of colonies 1, 2 and 3. Lane 1, 2 and 3 represent the gene of interest of 1287 bp. Lane M represents the standard markers.

#### Chikungunya virus rE2 gene expression

The pET28a(+)-rE2 plasmid was used to transform competent BL21(DE3)pLysS strain cells of *E. coli*. Before induction and after 16 h (overnight), samples were processed for protein extraction, and the expression profile of pET28a(+)-rE2 for purification on Ni-NTA column. Colony PCR confirmed the presence of rE2 gene (Figure 6C).

#### Expression and purification of rE2 in Nicotiana tabacum

The transgenic cassette containing the Chikungunya virus rE2 gene was successfully transferred from the pET28a(+) to pCAMBIA1302 (Supplementary Figure 8) at the multiple cloning site (MCS) (Figure 6A).

#### Mobilization of the recombinant vector pCAMBIA1302-CHIKV-E2 vector into A. tumefaciens (EHA105)

The recombinant vector pCAMBIA1302-rE2 was subsequently introduced into the virulent *Agrobacterium* EHA105 strain through the heat shock transformation method. Transformants were screened on YEM plates supplemented with 50 mg/mL kanamycin and 30 mg/mL rifampicin. White colonies were observed on the YEM agar plates (Figure 6B). The putative transformant clones exhibiting growth on the selective media were confirmed through colony PCR. Colony PCR was performed to verify the presence of the gene of interest in *Agrobacterium* (EHA105), and the results were analyzed on a 1.0% agarose gel (Figure 6C).

#### Transient transformation of the leaf of Nicotiana tabacum

Agrobacterium EHA105 with a unit absorption at OD_600_ concentration was infiltrated into the top elongated leaves of *Nicotiana tabacum* (Supplementary Figure 9), and leaf samples were collected at 3^rd^ and 8^th^ days post-infiltration (dpi). Transient transformation of the leaves was confirmed by confocal microscopy and Real-Time PCR.

#### Confocal microscopy confirmed CHIKV E2 expression in N. tabacum leaf

Under a confocal microscope, the transformed leaves of *N. tabacum* were validated by imaging. Visible expression of the green fluorescent protein (GFP) was evident 24 h post-transformation. The GFP expression was consistently observed on the third and eighth days of transformation. Maximum expression was visible at the 3^rd^-day post-infiltration (3^rd^-dpi) (Figure 7A-D).

**Figure 7.**
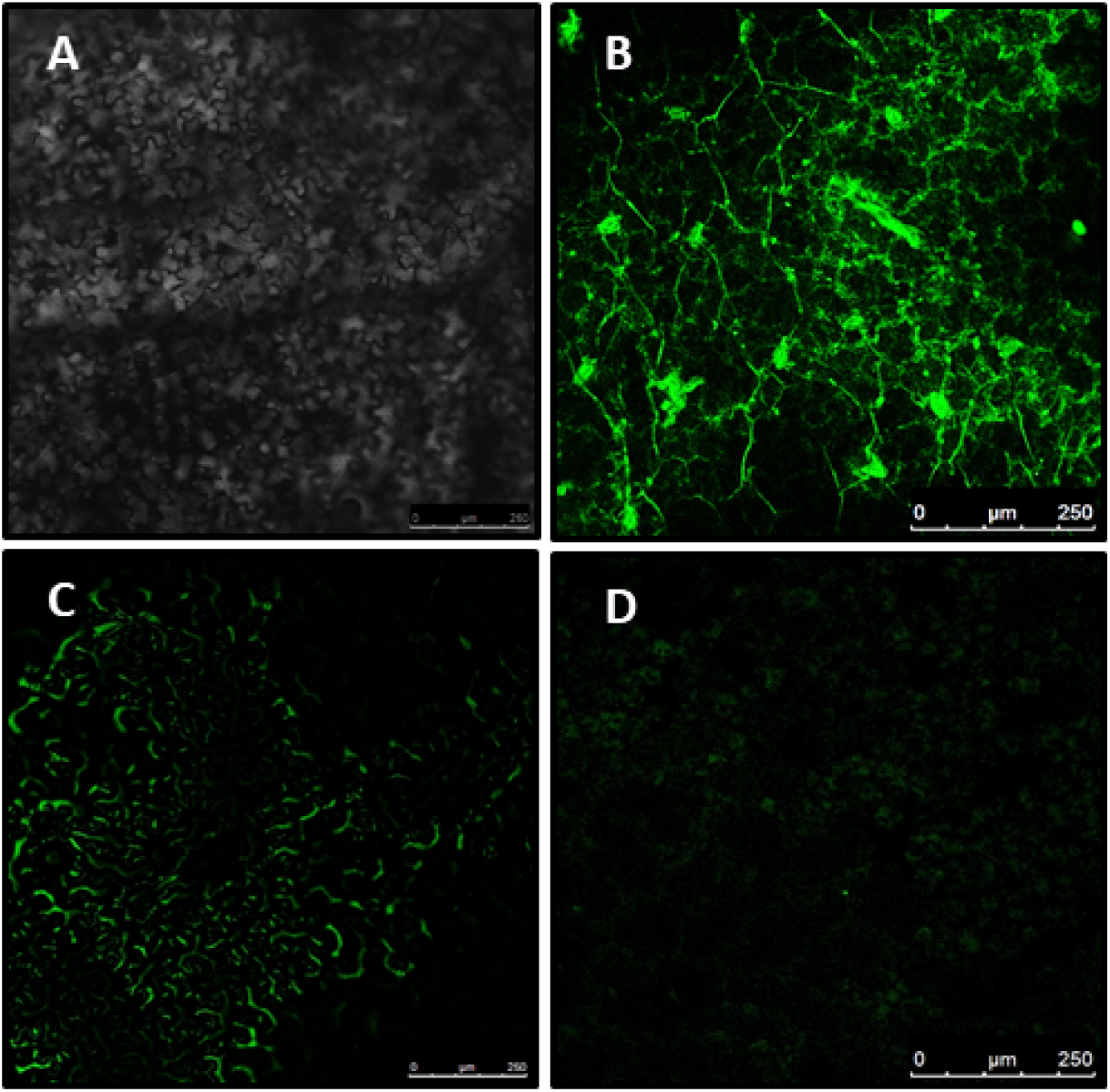
**(A-D)**: A: Control – Leaf of the wild-type tobacco (*Nicotiana tabacum*), B: GFP expression at 3^rd^-dpi, C: GFP expression at 8^th^-dpi, and D. expression at 15^th^-dpi.

#### Real-Time PCR of the CHIKV E2 in tobacco leaf

To confirm the molecular basis of confocal results, RT-PCR was performed with E2-specific primers. The amplified RT-PCR products yielded expected-sized amplicons on 1% agarose gel (Figure 8A). Quantitative expression estimation, with lower Cq values indicating higher transcript abundance in transiently transformed plants, confirmed the presence of E2 transcripts. Expression was more on the 3^rd^-dpi compared to the 8^th^-dpi. The ΔΔCt values for the control (wild-type plant) were zero. On the 3rd day post-infiltration, transgenic plant ΔΔCt values were 24.17, 23.22, and 23.71, with a mean value of 23.7. On the 8th day post-infiltration, ΔΔCt values were 21.76, 20.92, and 21.39, having a mean value of 21.3. Thus, the ΔΔCt value of the 3^rd^ dpi post-infiltration plant was higher than that of the 8^th^ dpi (Figure 8B).

**Figure 8.**
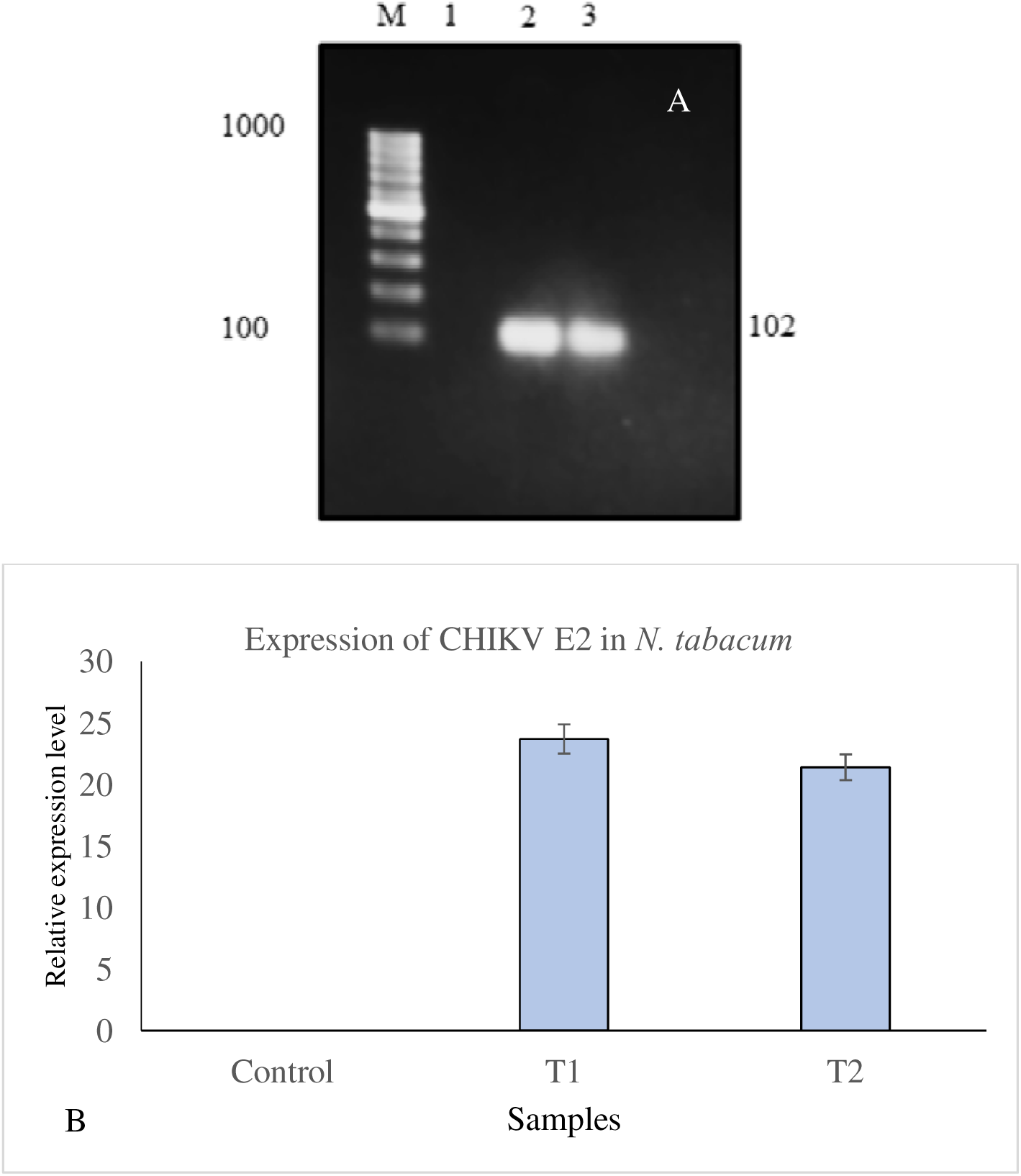
**(A-B): A.** RT-PCR product: Lane L contains a 100 bp ladder. Lane 1 contains the control sample of wild-type *N. tabacum*; Lane 2 contains a sample from a 3rd-day transient transformed sample, whereas Lane 3 contains an 8th-day sample from a transiently transformed *Nicotiana tabacum* plant. Bands of expected size were obtained on 1% agarose gel electrophoresis. **B.** The relative expression level (RT-PCR) for recombinant CHIKV E2 transcripts of the control and transiently transformed plants was estimated. No expression of recombinant E2 was found to be zero, whereas the expression level of the transiently transformed sample was highest for the 3^rd^ day (T1) sample as compared with the 8^th^ day (T2) sample.

#### Recombinant CHIKV rE2 protein from E. coli *and* N. tabacum had the same mass

Protein extracts of recombinant *E. coli* and transiently transformed tobacco were purified on a Ni-NTA affinity chromatography column. In both organisms, maximum yield was observed at 100 mM imidazole compared to 250 mM imidazole (Figure 9 A-C).

**Figure 9.**
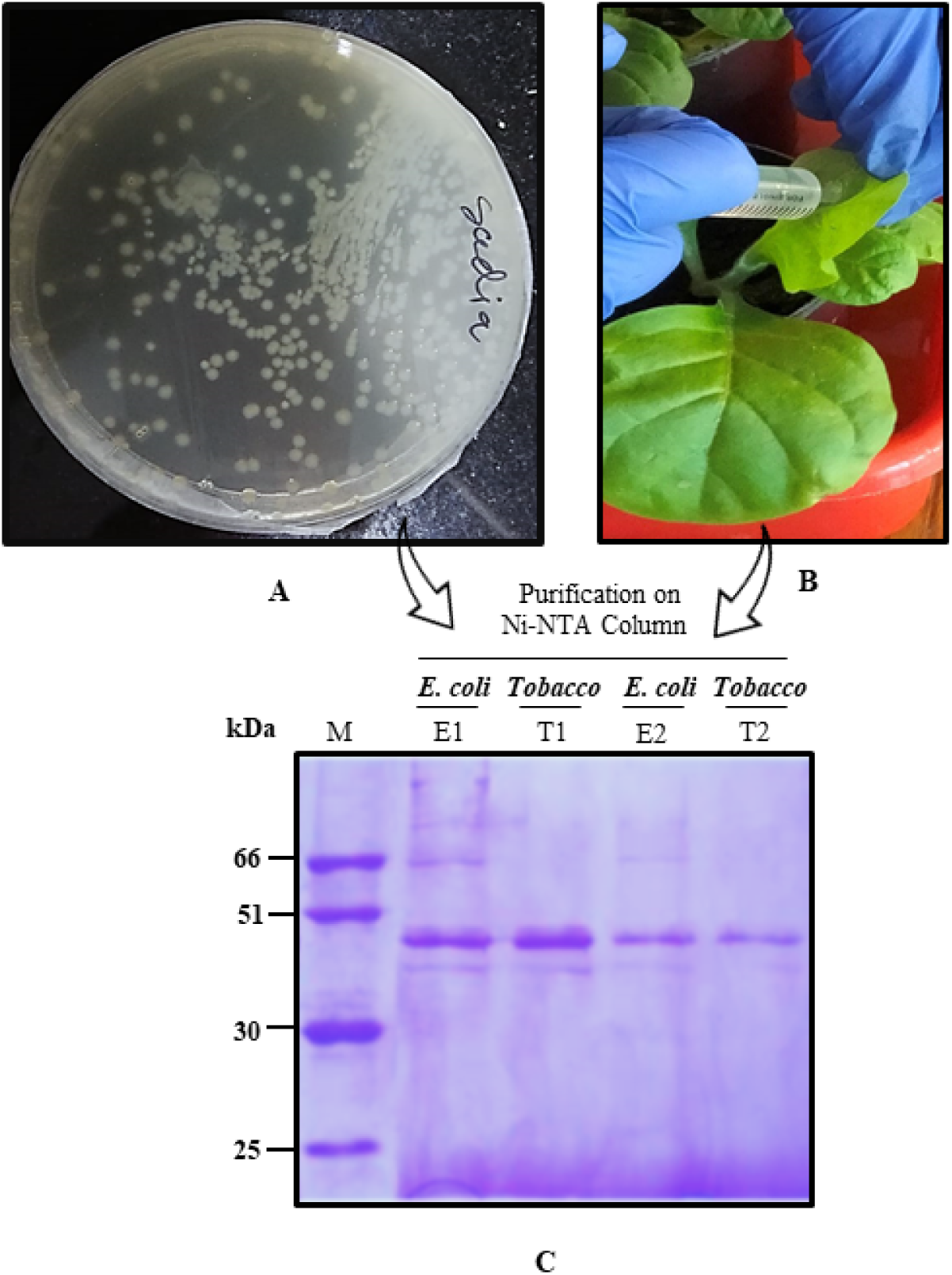
**(A-C): A.** Screening of genetically transformed *E. coli* to confirm cloning in BL21(DE3)pLysS strain cells. **B.** Transient expression of recombinant E2 through Agrobacterium-pCAMBIA-infiltration in the leaf of *N. tabacum.* **C.** Ni-NTA Affinity purified fractions from *E. coli* and *N. tabacum* loaded on 12% SDS-PAGE: Lane 1: Standard protein marker (M); Lane 2 (E1): Elution 1 (100 mM Imidazole) from *E. coli*, Lane 4 (E2): Elution 2 (250 mM Imidazole) from *E. coli*, Lane 4 (T1): Elution 3 (100 mM Imidazole) from *N. tabacum*, Lane 5 (T2): Elution 2 (250 mM Imidazole) from *N. tabacum* (Tobacco).

### Immunogenicity of CHIKV-rE2 antigen through indirect ELISA

Purified recombinant protein CHIKV E2 isolated from *E. coli* and *N. tabacum* was injected into BALB/c mice. Serum collected from both sets showed a strong presence of IgG in indirect ELISA (Figure 10 A-B). In both sets, a similar trend was observed. In an experiment with *E. coli*, immunization with rE2 protein + complete Freund’s adjuvant + incomplete Freund’s adjuvant (Group C) and rE2 protein + Alum (Group D) induced rE2-specific IgG antibodies, with a higher level observed in the CFA + IFA group compared to the Alum group. Recombinant CHIKV-E2 protein alone (Group B) also elicited IgG antibodies, though at a lower level than the adjuvanted groups. Control groups treated with normal saline showed no CHIKV-E2 IgG antibody responses, indicating the absence of pre-exposure to CHIKV-E2 antigen.

**Figure 10.**
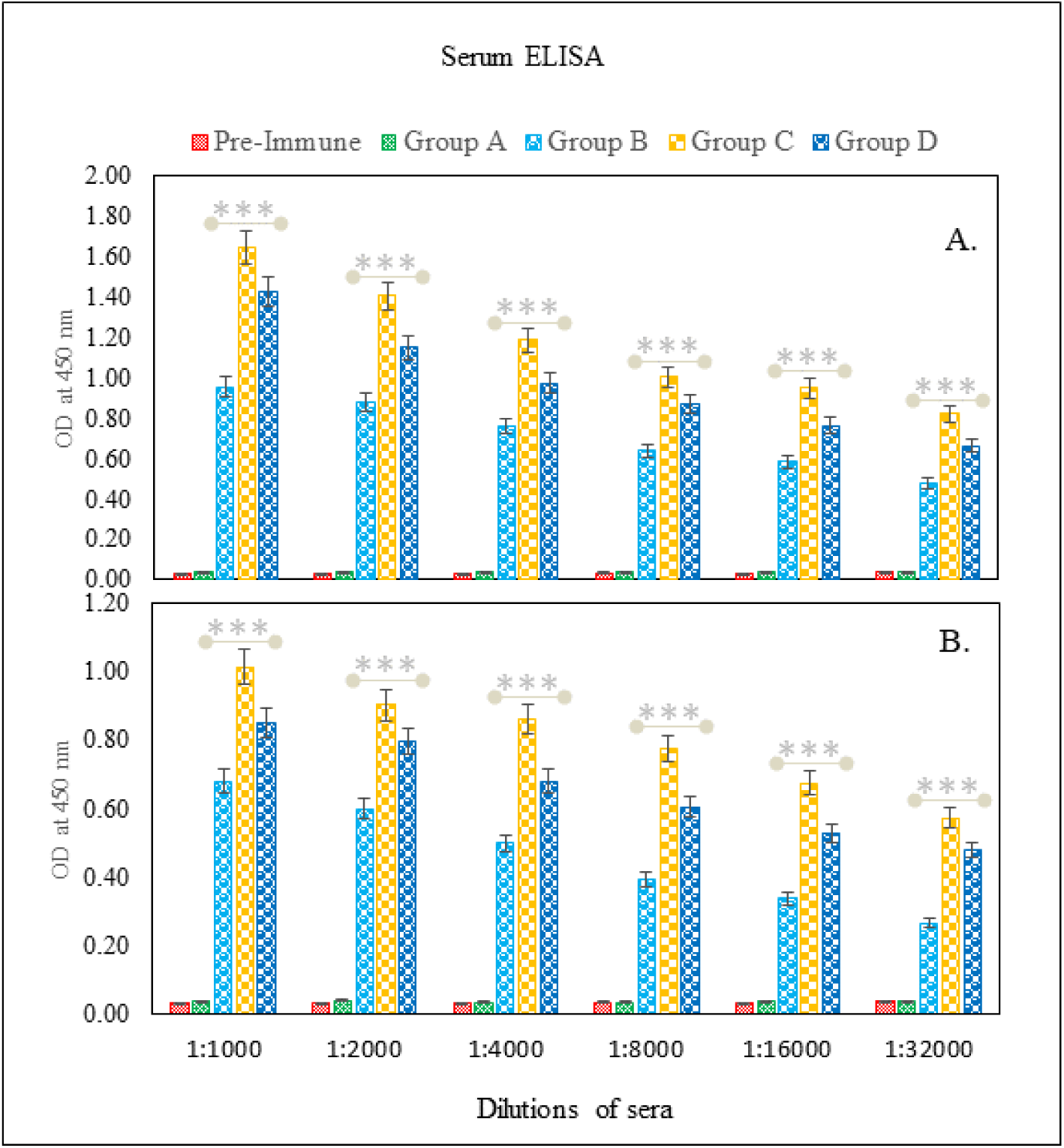
**(A-B):** ELISA of serum collected from BALB/c mice injected with recombinant E2 expressed in *E. coli* (A) and *N. tabacum* (B).

In a parallel experiment with *N. tabacum*, immunogenic responses were higher in Group C (rE2 protein from transgenic plant + CFA + IFA) compared to Group B (rE2 recombinant protein alone from transgenic plant) and Group D (rE2 protein from transgenic plant + Alum). Normalized titers, expressed as geometric mean titer (GMT) ± standard error (SE) per group, showed that the rE2 antigen inserted into both bacterial and plant systems was immunogenic, eliciting specific humoral responses. The results indicate the effectiveness of different immunization regimens and the potential of rE2 as an immunogen.

In brief, a positive result for rE2, expressed in *E. coli* (Figure 10A) or *N. tabacum* (Figure 10B) was obtained on the ELISA assay with highest antibody titer at a dilution of 1:1000. Importantly, Group C of BALB/c mice, which received recombinant E2 protein with complete Freund’s adjuvant and incomplete Freund’s adjuvant, exhibited the highest antibody titer and IgG response. The data were expressed as geometric mean titer (GMT) ± standard error (SE) per group using GraphPad Prism 5.0, highlighting the varying immunogenic responses in different treatment groups. One-way ANOVA was conducted, and the P value was found to be <0.0001 using Graphpad Prism-5. Tukey’s multiple comparison test was used to calculate the difference. The variance differs significantly, P <0.05, in all groups except Group A, as compared to the pre-immune sera. Group A was found to be non-significant when compared to the pre-immune sera.

### Western blot showed interaction of recombinant CHIKV E2 and BALB/c mice sera

Recombinant protein CHIKV E2, isolated from *E. coli* and tobacco (collected on the 3rd day post-infiltration), showed positive Western blots with nearly 49 kDa protein. The positive immunobinding of the rE2 protein with the developed antibody also confirmed the protein to be immunogenic (Figure 11).

**Figure 11:**
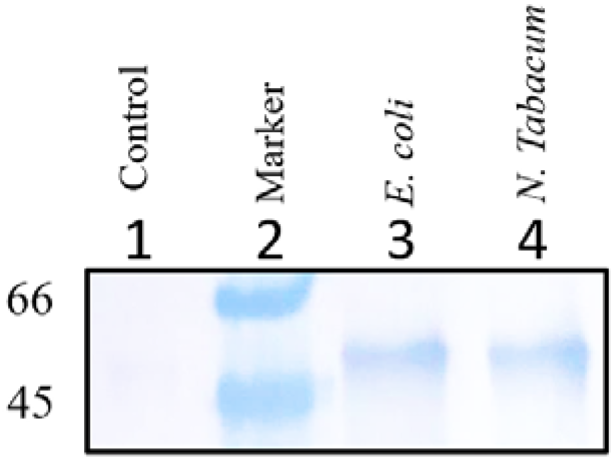
Western Blot image for recombinant protein; Lane 1: Control (sera from mice with no injection), Lane 2: Molecular marker; Lane 3: Purified antigen (10 μg from the mice injected with CHIKV E2 expressed in *E. coli*; and Lane 4: Purified antigen (10 μg from the mice injected with CHIKV E2 expressed in *N. tabacum*

## DISCUSSION

Our computational data predicted the success and efficacy of the CHIKV E2 spike protein, which plays a crucial role in the virus-host interaction (Shaikh et al., 2024), as a vaccine candidate. Importantly, there has been a notable gap in reports on the cloning and expression of the full-length E2 gene, particularly in the plant system, for vaccine development against Chikungunya infection. The merit of producing a vaccine candidate in a plant system lies in the ease of expression, high biomass yield, and, importantly, the large number of plant species that can serve as biofactories for vaccine production. In this study, we used the tobacco plant because of its various merits over other model plants. Tobacco is a preferred “green bioreactor” for producing recombinant proteins due to its high biomass yield, rapid growth, and cost-effective scalability compared to traditional systems like mammalian cell cultures. As a non-food crop, it reduces the risk of contaminating the food chain. Additionally, its well-established genetic transformation methods allow for the production of complex eukaryotic proteins with proper post-translational modifications (Naeem et al., 2024).

One of the features of CHIKV E2 is that it exhibits a very low frequency of Tryptophan, Phenylalanine, and Methionine. Tryptophan is the rarest, followed by Phenylalanine. This scarcity of large, bulky, and aromatic amino acids is common in many proteins, as they are energetically costly to produce and can impose significant structural constraints. The overall amino acid profile points to a protein with a robust, well-defined structure that incorporates both rigid and flexible elements, with a primary focus on hydrophobic interactions and specific folding patterns rather than a high prevalence of bulky aromatic residues (Supplementary Figure 1).

### Recombinant CHIKV E2 has significant possibilities for humanized PTMs

It had been a matter of concern whether the recombinant vaccine would work in the mammalian system or not. To address this, our results indicate a huge opportunity for the human body to humanize the recombinant protein at the residues that under undergoing PTMs as per human genome simulation rules (Figure 1 and Table 1). Thus, we were quite sure that the recombinant E2 vaccine had the potential to induce a humoral response in the BALB/c mice.

### In silico studies support E2 as a good vaccine candidate

We conducted *in silico* B-cell and T-cell epitope prediction for CHIKV E2-FL protein to comprehend immunogenicity, as B-cell and T-cell epitope content is one of the elements that affects protein antigenicity. Although there were some similar reports available (Chaudhary et al., 2024; Cho et al., 2024), we aimed to identify epitopes (Supplementary Figure 5 A-E) with updated portals/versions thoroughly (Supplementary Table 4 to Supplementary Table 9) and performed peptide-receptor molecular docking to assess the effectiveness (Figure 2A-E, Figure 3A-C, and Supplementary Table 10 to Supplementary Table 17). The NetCTL-1.2 server predicted five potent peptides, which showed a high conjunctional score from MHC-1. Furthermore, the study also identified four potent peptides from the SP protein of the Chikungunya virus for MHC Class-II potential peptide epitopes. The identified potent peptides had a high potential for the development of a vaccine against the Chikungunya virus. The peptide (YYYELYPTMTVVVVS) with the best IC50 value was evaluated using Molecular Dynamics (MD) Simulation (Figure 4A-E), showing impressive affinity.

### Molecular evaluation confirmed the putative transformants

After each step of the genetic transformation of *E. coli* and tobacco, molecular analysis was performed to assess the successful ligation or expression. Restriction/double digestion experiments confirmed the success of each step (Figure 5A-C). Success of the transformation of *A. tumefaciens* was evident from growth on selection media, followed by colony PCR (Figure 6A-C). Tobacco leaves with Agro-infiltration showed a transient expression of recombinant CHIKV E2, confirmed by confocal images. The high rate of expression on the 3^rd^ day of infiltration might be due to the optimal RNA-protein optimization dynamics, which later diminished with DNA-RNA-E2 degradation (Figure 7A-D). This data was supported by quantitative analysis of CHIKV E2 transcripts (Figure 8A-B). Isolation and purification of recombinant E2 on a Ni-NTA column, from both *E. coli* and tobacco, showed similar molecular weights (Figure 9).

### Immunogenicity of CHIKV-rE2 as an antigen

Numerous investigations have revealed that CHIKV envelope proteins can produce neutralizing antibodies that trigger a protective immune response, making them an attractive target for the CHIKV vaccine and diagnostic development (Khan et al., 2012; Islamuddin et al., 2022). Various studies on mice showed that an E2-based subunit vaccination was protective, and formulations of subunit vaccines based on the E2 protein induced a balanced Th1/Th2 response in mice and also developed virus-neutralizing antibodies. In the present study, the immunogenic potential of CHIKV E2 was estimated by immunizing the BALB/c mice with the recombinant spike protein E2 expressed in *E. coli* and *N. tabacum* along with CFA and IFA adjuvant and alum adjuvant. Analysis of antibody titers that are produced against the rE2 antigen in immunized mice using an Indirect ELISA test. We have shown that mice immunized with rE2 protein had a high titer of E2-specific antibodies. The immunogenicity assessment in BALB/c mice revealed higher IgG antibody levels for the rE2 protein + CFA + IFA combination compared to +Alum. A similar trend was observed in *N. tabacum*, emphasizing the potential of the CHIKV-E2 antigen in eliciting a humoral response.

In summary, various prokaryotic expression systems have been used in numerous investigations to report the cloning, production, and purification of E2 proteins of CHIKV E2 (Voss et al., 2010; Tripathi et al., 2014; Verma et al., 2016). In this study, we have performed cloning, expression, and purification of full-length CHIKV E2 protein in *E. coli* and *N. tabacum,* which not only expressed successfully but also induced humoral response in the mammalian model BALB/c mice.

## CONCLUSION

In summary, our study marks a significant step forward in providing a potent anti-chikungumya antigen/immunogen or vaccine candidate that has been successfully expressed in both bacteria and the plant system. We focused on cloning and expression of the full-length CHIKV E2 gene synthesized as per the S27 African prototype, a common chikungunya strain found in infected patients. As a major highlight, we explored *Agrobacterium*-mediated transient expression in *N. tabacum*, revealing dynamic expression levels, especially peaking on the 3^rd^ day post-infiltration. Our assessments of immunogenicity in BALB/c mice and *N. tabacum* showcased promising results, indicating the potential to elicit a strong immune response. This study not only enhances our understanding of CHIKV-host interactions but also paves the way for vaccine development and diagnostics. Our comprehensive approach positions us at the forefront of CHIKV studies, offering a holistic strategy to address Chikungunya virus infections. Moving forward, the insights gained from this study hold promise for further advancements, contributing to the global efforts to tackle Chikungunya virus infections.

## Acknowledgements

The authors express their sincere thanks to Jamia Millia Islamia (A Central University), New Delhi – 110025, India. First author is grateful to the University Grants Commission (UGC), GoI for providing Maulana Azad National Fellowship during her research.

