## Supplementary Material for "Spike protein E2 of chikungunya virus: a plant-based vaccine exhibited potent immunogenicity in BALB/c mice"

**Supplementary Table 1:** Nucleotide sequence of S27-African strain of CHIKV E2 gene

**CHIKV E2 envelope gene (GeneBank Accession No. AF339485.1)**

5’GGATCC (BamHI)

ATGAGCACCAAAGATAATTTCAATGTTTACAAAGCGACCCGTCCGTACCTGGCGCACTGCCCGGACTGCGGCGAAGGCCATAGCTGCCACAGCCCGGTGGCGCTGGAGCGTATCCGTAACGAAGCGACCGACGGCACCCTGAAGATTCAGGTTAGCCTGCAAATCGGTATTGGCACCGACGATAGCCACGATTGGACCAAACTGCGTTACATGGACAACCATATCCCGGCGGATGCGGGTCGTGCGGGCCTGTTTGTTCGTACCAGCGCGCCGTGCACCATCACCGGTACGATGGGTCACTTCATTCTGGCGCGTTGCCCGAAGGGTGAAACCCTGACCGTGGGCTTTACCGACAGCCGTAAAATCAGCCACAGCTGCACCCACCCGTTCCACCATGATCCGCCGGTTATTGGTCGTGAGAAGTTTCACAGCCGTCCGCAGCACGGCAAAGAACTGCCGTGCAGCACCTATGTGCAAAGCAACGCGGCGACCGCGGAGGAAATCGAAGTTCACATGCCGCCAGACACCCCGGATCGTACCCTGCTGAGCCAGCAAAGCGGTAACGTGAAGATTACCGTTAACGGCCGTACCGTTCGTTACAAATGCAACTGCGGTGGCAGCAACGAGGGTCTGATCACCACCGACAAAGTGATTAACAACTGCAAAGTTGATCAGTGCCACGCGGCGGTGACCAACCACAAGAAATGGCAATATAACAGCCCGCTGGTTCCGCGTAACGCGGAACTGGGTGACCGTAAGGGCAAAATCCACATTCCGTTCCCGCTGGCGAACGTGACCTGCATGGTTCCGAAGGCGCGTAACCCGACCGTGACCTACGGTAAAAACCAGGTTATCATGCTGCTGTATCCGGATCACCCGACCCTGCTGAGCTACCGTAGCATGGGCGAGGAACCGAACTATCAAGAGGAATGGGTGACCCACAAGAAAGAGGTGGTTCTGACCGTGCCGACCGAGGGTCTGGAAGTTACCTGGGGCAACAACGAACCGTATAAGTACTGGCCGCAGCTGAGCGCGAACGGCACCGCGCACGGCCACCCGCACGAGATCATTCTGTACTATTACGAACTGTACCCGACCATGACCGTGGTTGTGGTTAGCGTTGCGAGCTTTATCCTGCTGAGCATGGTGGGTATGGCGGTTGGCATGTGCATGTGCGCGCGTCGTCGTTGCATTACCCCGTATGAGCTGACCCCGGGTGCGACCGTTCCGTTCCTGCTGAGCCTGATTTGCTGCATCCGTACCGCGAAAGCGTAAAAGCTT HHHHHH3’ HindIII

**Supplementary Table 2:** Amino acid sequence of S27-African strain of CHIKV E2 protein

**CHIKV E2 envelope protein (Q8JUX5)**

**M**STKDNFNVYKATRPYLAHCPDCGEGHSCHSPVALERIRNEATDGTLKIQVSLQIGIGTDDSHDWTKLRYMDNHIPADAGRAGLFVRTSAPCTITGTMGHFILARCPKGETLTVGFTDSRKISHSCTHPFHHDPPVIGREKFHSRPQHGKELPCSTYVQSNAATAEEIEVHMPPDTPDRTLLSQQSGNVKITVNGRTVRYKCNCGGSNEGLITTDKVINNCKVDQCHAAVTNHKKWQYNSPLVPRNAELGDRKGKIHIPFPLANVTCMVPKARNPTVTYGKNQVIMLLYPDHPTLLSYRSMGEEPNYQEEWVTHKKEVVLTVPTEGLEVTWGNNEPYKYWPQLSANGTAHGHPHEIILYYYELYPTMTVVVVSVASFILLSMVGMAVGMCMCARRRCITPYELTPGATVPFLLSLICCIRTAKA**HHHHHH**

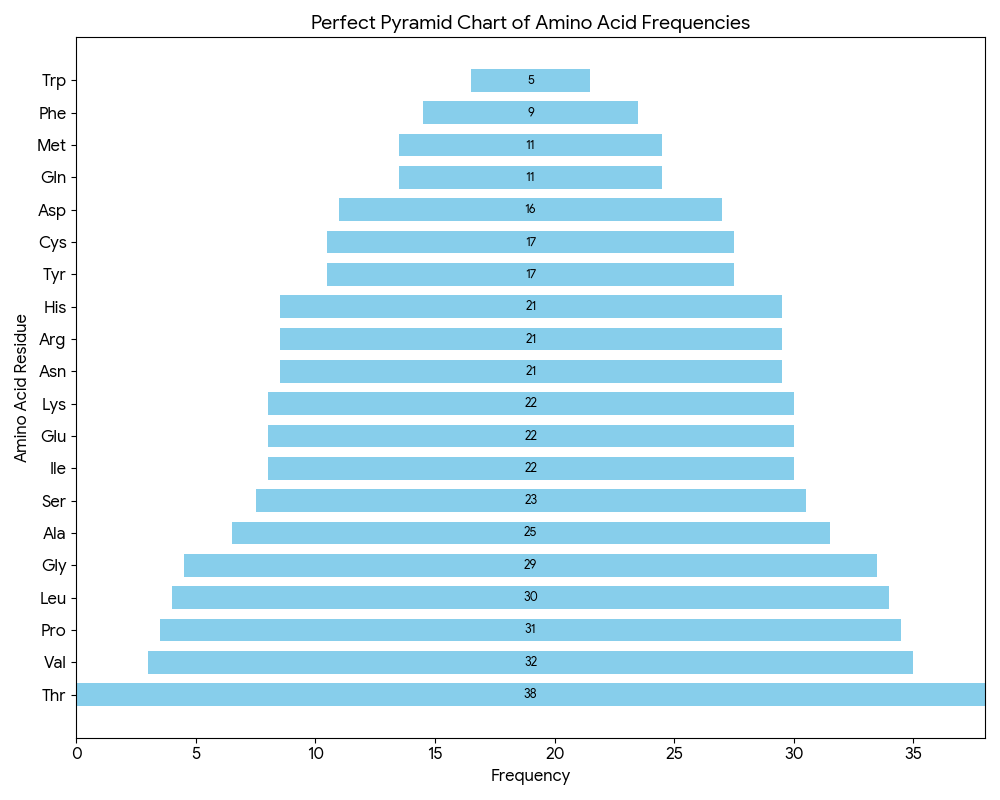

**Supplementary Figure 1:** Pyramid chart displaying the frequency of amino acid residues in a glimpse. There were only 5 residues of tryptophan (Trp) and 38 residues of threonine (Thr).

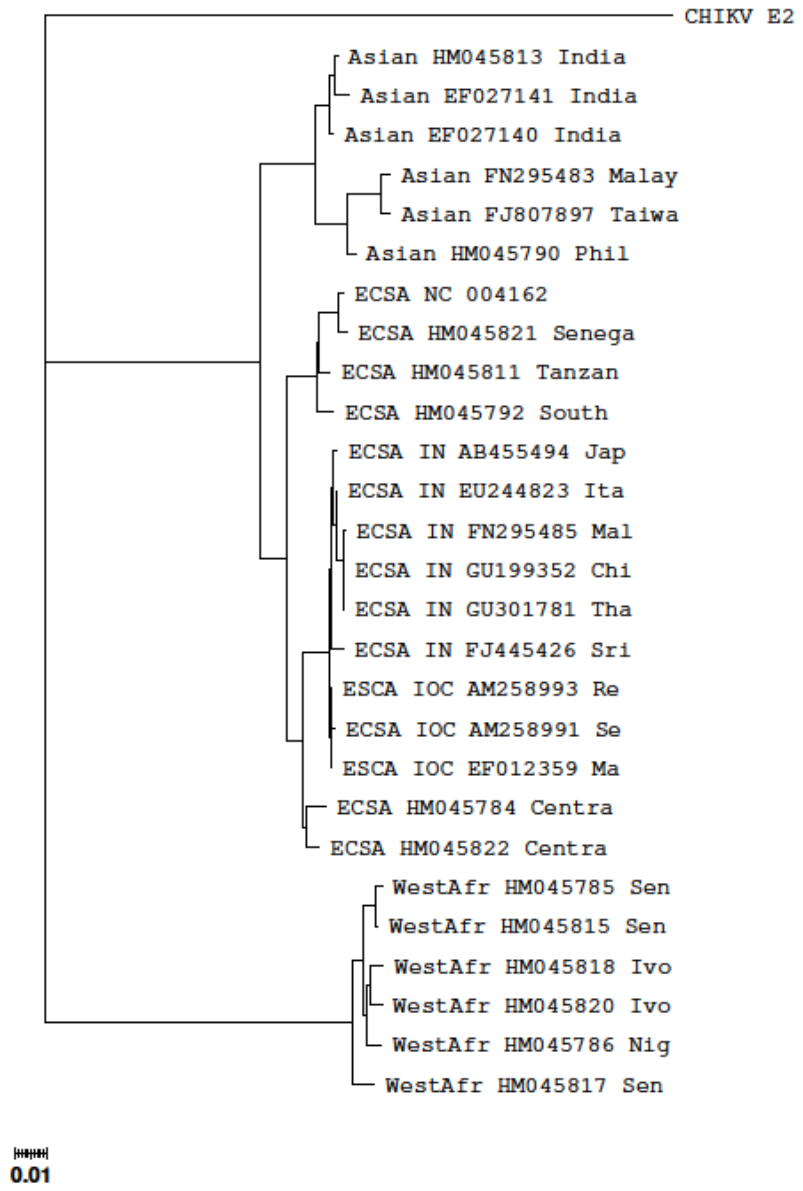

**Supplementary Figure 2:** Phylogenetic relationships among various serotypes of Chikungunya virus (Data obtained on 25 July 2025 from Chikungunya Typing Tool, Genome Detective) and the Clade assigned was ‘West African’. Supported with phylogenetic analysis and bootstrap 100.0 (>= 70.0). Genome region: Sequence starts at position 8539 and ends at position 9819 relative to the NC_004162.2 reference sequence for Chikungunya virus (taxon:37124).

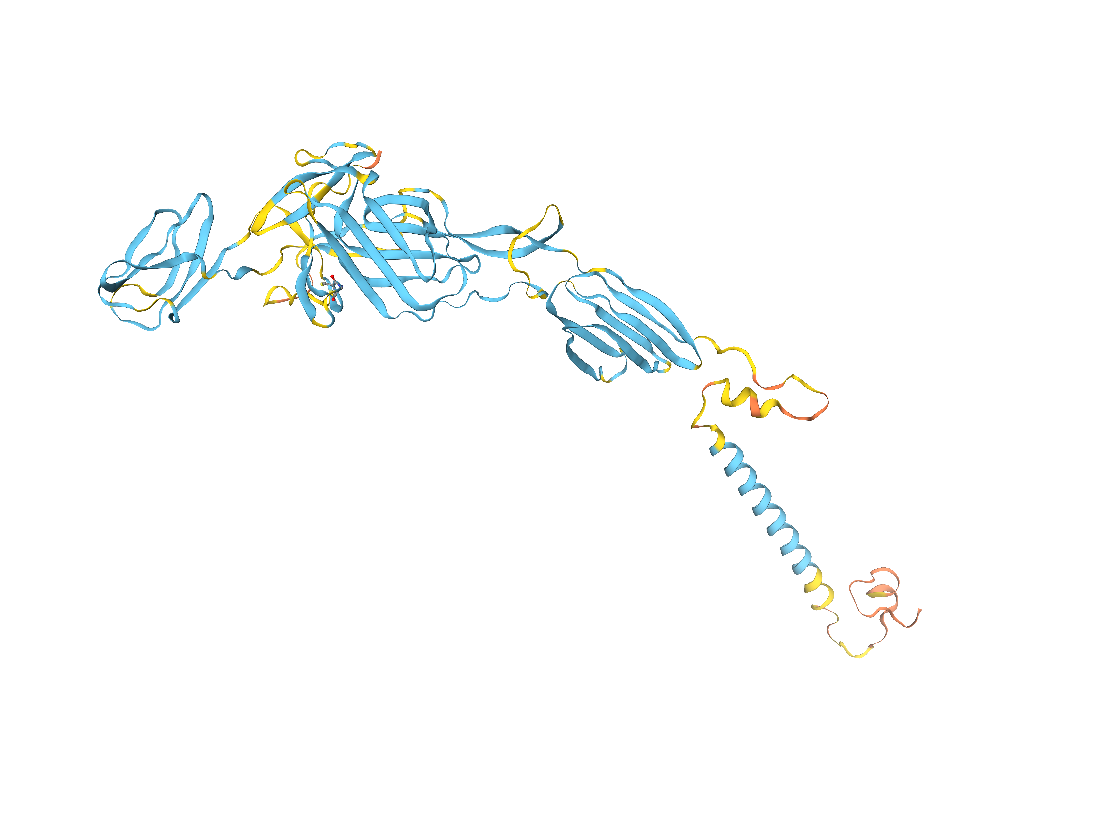

**Supplementary Figure 3:** A 3D model of CHIKV E2 generated on the SWISSDOCK portal, taking 6NK5 (PDB ID) as a template.

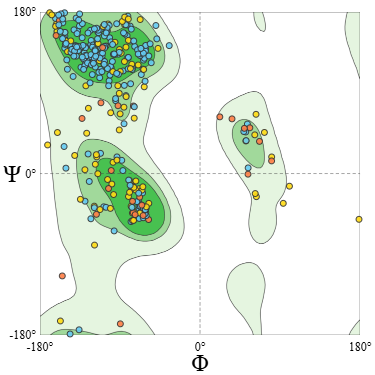

**Supplementary Figure 4:** The Ramachandran plot for CHIKV E2 proteins. Parameters and concerned values are explained in the Supplementary Table 2.

**Supplementary Table 3:** Multicrit table showing molprobity score, per cent of Ramachandran favored and outliers, rotamer outliers, C-beta deviations, bad bonds, bad angles, and twisted non-proline.

| MolProbity Score | 1.50 |
| --- | --- |
| Clash Score | 0.31 |
| Ramachandran Favoured | 88.49% |
| Ramachandran Outliers | 2.88% |
| n215 ASP, n174 PRO, n368 THR, n247 ALA, n355 GLU, n348 THR, n407 ALA, n410 PRO, n400 PRO, n366 THR, n403 LEU, n345 ALA | |
| Rotamer Outliers | 2.47% |
| n282 ASN, n209 GLU, n403 LEU, n106 CYS, n335 GLU, n412 LEU, n175 ASP, n267 CYS, n366 THR | |
| C-Beta Deviations | 21 |
| n364 TYR, n282 ASN, n61 ASP, n220 ASN, n366 THR, n414 SER, n379 LEU, n368 THR, n247 ALA, n25 GLU, n274 ASN, n186 SER, n121 LYS, n363 LEU, n399 THR, n372 VAL, n374 VAL, n376 SER, n418 CYS, n238 TYR, n74 HIS | |
| Bad Bonds | 0 / 3361 |
| Bad Angles | 44 / 4584 |
| (n399 THR-n400 PRO), (n404 THR-n405 PRO), (n240 SER-n241 PRO), n66 THR, n282 ASN, (n223 VAL-n224 ASP), n22 ASP, (n354 HIS-n355 GLU), (n219 ASN-n220 ASN), (n134 PRO-n135 PRO), n101 PHE, (n269 VAL-n270 PRO), n64 ASP, n233 HIS, n72 ASP, n124 HIS, n61 ASP, n30 HIS, (n352 HIS-n353 PRO), n8 ASN, n19 HIS, n352 HIS, n171 HIS, (n133 ASP-n134 PRO), (n172 MET-n173 PRO), (n152 LEU-n153 PRO), n131 HIS, n128 HIS, (n409 VAL-n410 PRO), n227 HIS, n231 THR, (n243 VAL-n244 PRO), n143 HIS, n132 HIS, n27 HIS, (n365 PRO-n366 THR), (n176 THR-n177 PRO), (n334 ASN-n335 GLU), (n128 HIS-n129 PRO), n93 THR, (n246 ASN-n247 ALA), n257 HIS, n63 HIS | |
| Twisted Non-Proline | 3 / 387 |
| (n23 CYS-n24 GLY), (n342 GLN-n343 LEU), (n343 LEU-n344 SER) | |

*Results obtained using MolProbity version 4.4*

**Supplementary Table 4:** B-cell epitope prediction using Emini Surface Accessibility Prediction (Predicted peptides) for CHIKV-E2 protein

| **S. No.** | **Start** | **End** | **Peptide** | **Length** |
| --- | --- | --- | --- | --- |
| 1 | 9 | 15 | YKATRPY | 7 |
| 2 | 36 | 42 | RIRNEAT | 7 |
| 3 | 59 | 70 | DDSHDWTKLRYM | 12 |
| 4 | 116 | 121 | TDSRKI | 6 |
| 5 | 139 | 149 | EKFHSRPQHGK | 11 |
| 6 | 172 | 178 | PPDTPDR | 7 |
| 7 | 230 | 238 | TNHKKWQYN | 9 |
| 8 | 248 | 253 | LGDRKG | 6 |
| 9 | 269 | 275 | PKARNPT | 7 |
| 10 | 297 | 315 | YRSMGEEPNYQEEWVTHKK | 19 |
| 11 | 331 | 340 | GNNEPYKYWP | 10 |

**Supplementary Table 5:** B-cell epitope prediction using Bepipred Linear Epitope Prediction 2.0 for CHIKV-E2 protein

| **S. No.** | **Start** | **End** | **Peptide** | **Length** |
| --- | --- | --- | --- | --- |
| 1 | 6 | 17 | FNVYKATRPYLA | 12 |
| 2 | 20 | 30 | PDCGEGHSCHS | 11 |
| 3 | 36 | 42 | RIRNEAT | 7 |
| 4 | 56 | 79 | IGTDDSHDWTKLRYMDNHIPADAG | 24 |
| 5 | 118 | 121 | SRKI | 4 |
| 6 | 131 | 167 | HDPPVIGREKFHSRPQHGKELPCSTYVQSNAATAEEI | 37 |
| 7 | 169 | 180 | VHMPPDTPDRTL | 12 |
| 8 | 207 | 212 | NEGLIT | 6 |
| 9 | 214 | 220 | DKVINNC | 7 |
| 10 | 234 | 252 | KWQYNSPLVPRNAELGDRK | 19 |
| 11 | 273 | 278 | NPTVTY | 6 |
| 12 | 300 | 314 | MGEEPNYQEEWVTHK | 15 |
| 13 | 330 | 350 | WGNNEPYKYWPQLSANGTAHG | 21 |

**Supplementary Table 6:** B-cell epitope prediction using BcePred (predicted B-cell epitopes) analysis for CHIKV-E2 protein using Chou & Fasman beta turn analysis and Parker Hydrophilicity prediction tool.

| **S.No.** | **Hydrophilicity** | **Flexibility** | **Accessibility** | **Turns** |
| --- | --- | --- | --- | --- |
| 1 | STKDNFN | TVGFTDSRK | STKDNFNVYKATRPYL | STKDNFNVY |
| 2 | CPDCGEGHSCHS | TLLSQQSGN | ALERIRNEATDGT | EGHSCHSPV |
| 3 | RNEATDGT | YKCNCGGSN | GTDDSHDWTKLRYMDNHI | GTDDSHDWT |
| 4 | GIGTDDSHDWT | AELGDRKG | RCPKGETLT | YMDNHIP |
| 5 | ADAGRAG | LSYRSMG | GFTDSRKISHS | ISHSCTHPFHHDPPV |
| 6 | RCPKGET |  | GREKFHSRPQHGKELP |  |
| 7 | SRPQHGKE |  | HMPPDTPDRTLLSQQSGNVK |  |
| 8 | SNAATAEE |  | NGRTVRYKCN |  |
| 9 | PPDTPDRT |  | NNCKVDQ |  |
| 10 | SQQSGNVK |  | AVTNHKKWQYNSP |  |
| 11 | KCNCGGSNEG |  | AELGDRKGKIH |  |
| 12 | NNCKVDQ |  | MVPKARNPTVTYGKNQV |  |
| 13 | SMGEEPNYQEE |  | LYPDHPTL |  |
| 14 | GNNEPYK |  | SYRSMGEEPNYQEEWVTHKKEVV |  |
| 15 |  |  | WGNNEPYKYWPQLS |  |
| 16 |  |  | YYYELYPT |  |
| 17 |  |  | RRRCITPYELTP |  |

**Supplementary Table 7:** Kolaskar & Tongaonkar Antigenicity (Predicted peptides) for CHIKV-E2 protein

| **S.No.** | **Start** | **End** | **Peptide** | **Length** |
| --- | --- | --- | --- | --- |
| 1 | 14 | 22 | PYLAHCPDC | 9 |
| 2 | 27 | 35 | SCHSPVALE | 9 |
| 3 | 47 | 54 | KIQVSLQI | 8 |
| 4 | 82 | 88 | GLFVRTS | 7 |
| 5 | 100 | 105 | FILARC | 6 |
| 6 | 122 | 128 | SHSCTHP | 7 |
| 7 | 130 | 135 | HHDPPV | 6 |
| 8 | 150 | 156 | ELPCSTY | 7 |
| 9 | 198 | 203 | RYKCNC | 6 |
| 10 | 222 | 229 | VDQCHAAV | 8 |
| 11 | 256 | 269 | HIPFPLANVTCMVP | 14 |
| 12 | 283 | 297 | VIMLLYPDHPTLLSY | 15 |
| 13 | 314 | 321 | KKEVVLTV | 8 |
| 14 | 353 | 362 | HEIILYYYEL | 10 |
| 15 | 365 | 381 | TMTVVVVSVASFILLSM | 17 |
| 16 | 388 | 394 | MCMCARR | 7 |

**Supplementary Table 8:** T-cell epitopes prediction for CHIKV E2. Five potent CD8+ T-cell epitopes and their interacting MHC Class-I alleles and NetCTL combine score, immunogenicity score and epitope conservancy hits.

| **Peptide start** | **Peptide end** | **Peptide** | **IC50** | **TepiTool result Allele** | **NetCTL combined score** | **Immunogenicity** |
| --- | --- | --- | --- | --- | --- | --- |
| 351 | 359 | HPHEIILYY | 5.6  29.6 | HLA-B*35:01;  HLA-B*53:01 | A1- 0.7537; A3- 0.7833 | 0.35461 |
| 298 | 306 | RSMGEEPNY | 56.7  58.7 | HLA-B*58:01  HLA-A*30:02 | A1- 0.8313; A3- 0.7767; B58- 1.7764; B62- 1.1900 | 0.17163 |
| 280 | 288 | KNQVIMLLY | 57.6 | HLA-A*30:02 | A1- 0.8541 | 0.15571 |
| 1 | 9 | STKDNFNVY | 65.3  81.2 | HLA-A*30:02  HLA-B*15:01 | A1- 2.1435; A26- 2.3492; B62- 1.3092 | 0.07488 |
| 328 | 336 | VTWGNNEPY | 97.9 | HLA-B*35:01 | A1- 1.2063; A26- 1.5023; B58- 0.7522; B62- 1.3173 | -0.0476 |

**Supplementary Table 9**: The selected four most potent CD4+ T-cell epitopes and their interacting MHC class-II alleles with affinity IC50 < 100.

| Peptide start | Peptide end | Peptide sequence | IC50 | Allele |
| --- | --- | --- | --- | --- |
| 4 | 18 | DNFNVYKATRPYLAH | 14.5 | HLA-DRB1*01:01 |
|  |  |  | 27.55 | HLA-DRB1*07:01 |
|  |  |  | 28.24 | HLA-DRB5*01:01 |
|  |  |  | 39.08 | HLA-DRB1*09:01 |
|  |  |  | 49.72 | HLA-DRB1*11:01 |
|  |  |  | 79.41 | HLA-DRB1*15:01 |
| 80 | 94 | RAGLFVRTSAPCTIT | 17.35 | HLA-DRB1*01:01 |
|  |  |  | 37.23 | HLA-DRB1*07:01 |
|  |  |  | 49.03 | HLA-DRB1*09:01 |
|  |  |  | 62.33 | HLA-DRB3*02:02 |
|  |  |  | 91.38 | HLA-DRB1*04:01 |
|  |  |  | 92.43 | HLA-DRB1*13:02 |
| 358 | 372 | YYYELYPTMTVVVVS | 17.47 | HLA-DRB1*01:01 |
|  |  |  | 25.93 | HLA-DRB1*07:01 |
|  |  |  | 45.92 | HLA-DRB1*09:01 |
|  |  |  | 64.52 | HLA-DRB1*13:02 |

**Supplementary Table 10**: Interaction of predicted CD8+ T-cell epitope “HPHEIILYY” in CHIKV E2 with MHC class-I alleles having varied affinity of IC50.

| **Name** | **Distance** | **Category** | **Type** |
| --- | --- | --- | --- |
| A:ARG65:HH12 - B:GLU278:OE1 | 2.06045 | Hydrogen Bond;Electrostatic | Salt Bridge;Attractive Charge |
| A:LYS146:HZ1 - B:TYR283:OXT | 1.92729 | Hydrogen Bond;Electrostatic | Salt Bridge;Attractive Charge |
| B:HIS275:HT3 - A:GLU63:OE2 | 1.91927 | Hydrogen Bond;Electrostatic | Salt Bridge;Attractive Charge |
| A:ASN66:HD22 - B:PRO276:O | 2.10492 | Hydrogen Bond | Conventional H-Bond |
| A:TYR99:HH - B:HIS277:O | 2.71376 | Hydrogen Bond | Conventional H-Bond |
| A:LYS146:HZ3 - B:TYR282:OH | 2.76326 | Hydrogen Bond | Conventional H-Bond |
| A:TRP147:HE1 - B:LEU281:O | 2.79356 | Hydrogen Bond | Conventional H-Bond |
| A:TYR159:HH - B:HIS275:O | 2.02814 | Hydrogen Bond | Conventional H-Bond |
| A:TRP167:HE1 - B:HIS275:O | 2.2601 | Hydrogen Bond | Conventional H-Bond |
| B:HIS275:HT3 - A:TYR59:OH | 2.05184 | Hydrogen Bond | Conventional H-Bond |
| B:HIS277:HN - A:TYR99:OH | 2.45756 | Hydrogen Bond | Conventional H-Bond |
| B:ILE279:HN - B:HIS277:O | 2.97462 | Hydrogen Bond | Conventional H-Bond |
| B:TYR282:HN - A:ASP77:OD2 | 2.06365 | Hydrogen Bond | Conventional H-Bond |
| B:HIS275:CE1 - A:GLN62:OE1 | 3.69996 | Hydrogen Bond | Carbon H-Bond |
| B:PRO276:CD - A:GLU63:OE1 | 3.27991 | Hydrogen Bond | Carbon H-Bond |
| B:HIS275:HT1 - A:TRP167 | 2.35084 | Hydrogen Bond;Electrostatic | Pi-Cation;Pi-Donor Hydrogen Bond |
| B:HIS275:HT1 - A:TRP167 | 3.17661 | Hydrogen Bond;Electrostatic | Pi-Cation;Pi-Donor Hydrogen Bond |
| B:TYR283:OXT - B:TYR282 | 3.36397 | Electrostatic | Pi-Anion |
| B:TYR282:HH - A:TYR84 | 3.05578 | Hydrogen Bond | Pi-Donor Hydrogen Bond |
| B:HIS277:CB - A:TYR159 | 3.77354 | Hydrophobic | Pi-Sigma |
| B:PRO276:C,O;HIS277:N - A:TYR159 | 4.40198 | Hydrophobic | Amide-Pi Stacked |
| A:VAL67 - B:PRO276 | 5.3872 | Hydrophobic | Alkyl |
| A:VAL76 - B:LEU281 | 5.05602 | Hydrophobic | Alkyl |
| A:ILE97 - B:ILE279 | 5.08562 | Hydrophobic | Alkyl |
| A:ALA150 - B:ILE280 | 4.66866 | Hydrophobic | Alkyl |
| A:ALA152 - B:ILE280 | 4.45087 | Hydrophobic | Alkyl |
| B:TYR282 - A:LYS146 | 5.16563 | Hydrophobic | Pi-Alkyl |
| B:TYR283 - A:LYS146 | 5.21942 | Hydrophobic | Pi-Alkyl |

**Supplementary Table 11:** Interaction of predicted CD8+ T-cell epitope “RSMGEEPNY” in CHIKV E2 with MHC class-I alleles having varied affinity of IC50.

| **Name** | **Distance** | **Category** | **Type** |
| --- | --- | --- | --- |
| B:ARG275:HH21 - A:GLU63:OE2 | 1.98041 | Hydrogen Bond;Electrostatic | Salt Bridge;Attractive Charge |
| B:ARG275:N - A:GLU63:OE2 | 5.14566 | Electrostatic | Attractive Charge |
| A:ASN66:HD22 - B:ARG275:O | 1.98317 | Hydrogen Bond | Conventional H-Bond |
| A:GLN70:HE21 - B:SER276:OG | 2.30464 | Hydrogen Bond | Conventional H-Bond |
| A:GLN70:HE22 - B:SER276:OG | 2.49995 | Hydrogen Bond | Conventional H-Bond |
| A:TYR99:HH - B:SER276:OG | 2.53794 | Hydrogen Bond | Conventional H-Bond |
| A:GLN155:HE22 - B:GLU279:OE1 | 2.00423 | Hydrogen Bond | Conventional H-Bond |
| A:TYR159:HH - B:SER276:O | 2.00717 | Hydrogen Bond | Conventional H-Bond |
| A:TRP167:HE1 - B:SER276:O | 2.72412 | Hydrogen Bond | Conventional H-Bond |
| B:ARG275:HE - A:GLU63:OE2 | 1.8715 | Hydrogen Bond | Conventional H-Bond |
| B:ARG275:HH22 - A:TYR7:OH | 2.171 | Hydrogen Bond | Conventional H-Bond |
| B:ARG275:HH22 - A:TYR171:OH | 2.17588 | Hydrogen Bond | Conventional H-Bond |
| B:SER276:HN - A:GLU63:OE2 | 2.1137 | Hydrogen Bond | Conventional H-Bond |
| B:SER276:HG - A:TYR9:OH | 2.02855 | Hydrogen Bond | Conventional H-Bond |
| B:GLY278:HN - B:ARG275:O | 2.15676 | Hydrogen Bond | Conventional H-Bond |
| B:GLU279:HN - B:MET277:O | 2.88267 | Hydrogen Bond | Conventional H-Bond |
| B:ASN282:HN - A:GLN155:OE1 | 2.74994 | Hydrogen Bond | Conventional H-Bond |
| B:ASN282:HD21 - A:GLN155:OE1 | 2.75671 | Hydrogen Bond | Conventional H-Bond |
| B:PRO281:CA - A:GLN155:OE1 | 3.71325 | Hydrogen Bond | Carbon H-Bond |
| B:ARG275:NH1 - A:TYR171 | 4.85017 | Electrostatic | Pi-Cation |
| B:ARG275:NH2 - A:TRP167 | 4.73482 | Electrostatic | Pi-Cation |
| A:ASP77:OD2 - B:TYR283 | 3.7123 | Electrostatic | Pi-Anion |
| B:TYR283:OXT - A:TRP147 | 4.42269 | Electrostatic | Pi-Anion |
| B:ARG275:CD - A:TRP167 | 3.95867 | Hydrophobic | Pi-Sigma |
| B:MET277:SD - A:TYR99 | 3.61411 | Other | Pi-Sulfur |
| A:TRP147 - B:TYR283 | 5.2318 | Hydrophobic | Pi-Pi T-shaped |
| B:ARG275:C,O;SER276:N - A:TRP167 | 5.00522 | Hydrophobic | Amide-Pi Stacked |
| B:ARG275:C,O;SER276:N - A:TRP167 | 5.15074 | Hydrophobic | Amide-Pi Stacked |
| B:PRO281 - B:MET277 | 4.24989 | Hydrophobic | Alkyl |
| A:TYR59 - B:ARG275 | 5.24946 | Hydrophobic | Pi-Alkyl |

**Supplementary Table 12:** Interaction of predicted CD8+ T-cell epitope “KNQVIMLLY” in CHIKV E2 with MHC class-I alleles having varied affinity of IC50.

| **Name** | **Distance** | **Category** | **Type** |
| --- | --- | --- | --- |
| A:LYS146:HZ1 - B:TYR283:OXT | 2.37599 | Hydrogen Bond;Electrostatic | Salt Bridge;Attractive Charge |
| B:LYS275:N - A:GLU63:OE2 | 4.30135 | Electrostatic | Attractive Charge |
| B:LYS275:NZ - A:GLU63:OE2 | 5.27549 | Electrostatic | Attractive Charge |
| A:TYR9:HH - B:ASN276:OD1 | 2.41521 | Hydrogen Bond | Conventional H-Bond |
| A:TYR84:HH - B:TYR283:OXT | 2.14484 | Hydrogen Bond | Conventional H-Bond |
| A:TYR99:HH - B:LYS275:O | 2.81295 | Hydrogen Bond | Conventional H-Bond |
| A:TRP147:HE1 - B:LEU281:O | 2.88743 | Hydrogen Bond | Conventional H-Bond |
| A:TRP147:HE1 - B:LEU282:O | 2.47205 | Hydrogen Bond | Conventional H-Bond |
| A:TYR159:HH - B:LYS275:O | 2.03846 | Hydrogen Bond | Conventional H-Bond |
| B:LYS275:HT1 - A:TYR171:OH | 2.64739 | Hydrogen Bond | Conventional H-Bond |
| B:LYS275:HT3 - A:TYR59:OH | 2.07487 | Hydrogen Bond | Conventional H-Bond |
| B:LYS275:HT3 - A:TYR171:OH | 2.9023 | Hydrogen Bond | Conventional H-Bond |
| B:LYS275:HZ1 - A:GLN62:OE1 | 2.69565 | Hydrogen Bond | Conventional H-Bond |
| B:LYS275:HZ3 - A:GLN62:OE1 | 2.2526 | Hydrogen Bond | Conventional H-Bond |
| B:ASN276:HN - A:GLU63:OE2 | 2.5014 | Hydrogen Bond | Conventional H-Bond |
| B:ASN276:HD22 - A:GLU63:O | 2.25495 | Hydrogen Bond | Conventional H-Bond |
| B:GLN277:HN - A:TYR99:OH | 2.28826 | Hydrogen Bond | Conventional H-Bond |
| B:VAL278:HN - B:GLN277:OE1 | 2.51507 | Hydrogen Bond | Conventional H-Bond |
| B:LYS275:CA - A:GLU63:OE2 | 3.27869 | Hydrogen Bond | Carbon H-Bond |
| B:LEU282:CA - A:ASP77:OD2 | 3.54445 | Hydrogen Bond | Carbon H-Bond |
| B:LYS275:N - A:TRP167 | 3.93581 | Electrostatic | Pi-Cation |
| B:LYS275:CB - A:TRP167 | 3.82013 | Hydrophobic | Pi-Sigma |
| A:TRP147 - B:TYR283 | 4.82449 | Hydrophobic | Pi-Pi T-shaped |
| A:ALA69 - B:ILE279 | 3.8478 | Hydrophobic | Alkyl |
| A:VAL76 - B:LEU282 | 5.45773 | Hydrophobic | Alkyl |
| A:ALA150 - B:LEU281 | 4.92183 | Hydrophobic | Alkyl |
| A:ALA152 - B:MET280 | 5.46019 | Hydrophobic | Alkyl |
| A:TYR59 - B:LYS275 | 5.38189 | Hydrophobic | Pi-Alkyl |
| A:TRP147 - B:MET280 | 5.07378 | Hydrophobic | Pi-Alkyl |
| A:TRP167 - B:LYS275 | 5.34097 | Hydrophobic | Pi-Alkyl |
| B:TYR283 - A:LEU81 | 5.37645 | Hydrophobic | Pi-Alkyl |

**Supplementary Table 13:**: Interaction of predicted CD8+ T-cell epitope “STKDNFNVY” in CHIKV E2 with MHC class-I alleles having varied affinity of IC50.

| **Name** | **Distance** | **Category** | **Type** |
| --- | --- | --- | --- |
| A:ARG65:HH21 - B:ASP278:OD1 | 3.18492 | Hydrogen Bond;Electrostatic | Salt Bridge;Attractive Charge |
| A:ARG65:HH22 - B:ASP278:OD1 | 3.19623 | Hydrogen Bond;Electrostatic | Salt Bridge;Attractive Charge |
| A:ARG163:HH21 - B:ASP278:OD2 | 3.0311 | Hydrogen Bond;Electrostatic | Salt Bridge;Attractive Charge |
| B:SER275:N - A:GLU63:OE2 | 3.97293 | Electrostatic | Attractive Charge |
| A:TYR59:HH - B:SER275:OG | 2.35628 | Hydrogen Bond | Conventional H-Bond |
| A:ASN66:HD21 - B:ASP278:OD1 | 1.99884 | Hydrogen Bond | Conventional H-Bond |
| A:ASN66:HD22 - B:THR276:OG1 | 2.50095 | Hydrogen Bond | Conventional H-Bond |
| A:ASN66:HD22 - B:THR276:O | 2.33955 | Hydrogen Bond | Conventional H-Bond |
| A:GLN70:HE21 - B:ASN281:OD1 | 2.41907 | Hydrogen Bond | Conventional H-Bond |
| A:TYR99:HH - B:SER275:O | 2.68928 | Hydrogen Bond | Conventional H-Bond |
| A:TYR159:HH - B:ASP278:OD2 | 2.13807 | Hydrogen Bond | Conventional H-Bond |
| A:ARG163:HH12 - B:ASN279:OD1 | 2.0172 | Hydrogen Bond | Conventional H-Bond |
| A:ARG163:HH22 - B:ASN279:OD1 | 2.67687 | Hydrogen Bond | Conventional H-Bond |
| B:SER275:HT3 - A:TYR7:OH | 1.88156 | Hydrogen Bond | Conventional H-Bond |
| B:SER275:HT3 - A:TYR171:OH | 2.31467 | Hydrogen Bond | Conventional H-Bond |
| B:THR276:HN - A:GLU63:OE2 | 2.723 | Hydrogen Bond | Conventional H-Bond |
| B:THR276:HG1 - A:GLU63:O | 2.32215 | Hydrogen Bond | Conventional H-Bond |
| B:LYS277:HN - A:TYR99:OH | 2.19602 | Hydrogen Bond | Conventional H-Bond |
| B:LYS277:HZ1 - B:ASN279:O | 1.89673 | Hydrogen Bond | Conventional H-Bond |
| B:LYS277:HZ3 - B:ASN281:OD1 | 1.89424 | Hydrogen Bond | Conventional H-Bond |
| B:ASN279:HD21 - A:GLN155:O | 2.81061 | Hydrogen Bond | Conventional H-Bond |
| B:ASN279:HD22 - A:GLN155:O | 2.52748 | Hydrogen Bond | Conventional H-Bond |
| B:ASN281:HD21 - A:GLN70:OE1 | 2.05425 | Hydrogen Bond | Conventional H-Bond |
| A:TRP167:CD1 - B:SER275:O | 3.52837 | Hydrogen Bond | Carbon H-Bond |
| B:SER275:CA - A:GLU63:OE2 | 3.11728 | Hydrogen Bond | Carbon H-Bond |
| B:SER275:HT2 - A:TRP167 | 2.64604 | Hydrogen Bond;Electrostatic | Pi-Cation;Pi-Donor H-Bond |
| A:ALA150 - B:VAL282 | 3.68598 | Hydrophobic | Alkyl |
| A:ALA152 - B:VAL282 | 4.94269 | Hydrophobic | Alkyl |
| A:TYR99 - B:LYS277 | 4.90646 | Hydrophobic | Pi-Alkyl |
| A:TRP147 - B:VAL282 | 5.14877 | Hydrophobic | Pi-Alkyl |
| A:TYR159 - B:LYS277 | 4.85832 | Hydrophobic | Pi-Alkyl |
| B:PHE280 - A:ALA69 | 5.02337 | Hydrophobic | Pi-Alkyl |

**Supplementary Table 14:** Interaction of predicted CD8+ T-cell epitope “VTWGNNEPY” in CHIKV E2 with MHC class-I alleles having varied affinity of IC50.

| **Name** | **Distance** | **Category** | **Type** |
| --- | --- | --- | --- |
| A:ARG114:HH12 - B:GLU281:OE1 | 1.83585 | Hydrogen Bond;Electrostatic | Salt Bridge;Attractive Charge |
| A:ARG114:HH22 - B:GLU281:OE1 | 2.81755 | Hydrogen Bond;Electrostatic | Salt Bridge;Attractive Charge |
| B:VAL275:HT1 - A:GLU63:OE1 | 1.97514 | Hydrogen Bond;Electrostatic | Salt Bridge;Attractive Charge |
| B:VAL275:HT2 - A:GLU63:OE2 | 2.25252 | Hydrogen Bond;Electrostatic | Salt Bridge;Attractive Charge |
| A:ARG114:NH2 - B:GLU281:OE2 | 4.79647 | Electrostatic | Attractive Charge |
| A:LYS146:NZ - B:TYR283:OXT | 4.78825 | Electrostatic | Attractive Charge |
| A:ARG65:HH12 - B:GLY278:O | 2.03986 | Hydrogen Bond | Conventional H-Bond |
| A:ARG65:HH22 - B:GLY278:O | 2.36498 | Hydrogen Bond | Conventional H-Bond |
| A:TYR84:HH - B:TYR283:OH | 2.70831 | Hydrogen Bond | Conventional H-Bond |
| A:TYR159:HH - B:VAL275:O | 2.08764 | Hydrogen Bond | Conventional H-Bond |
| A:TRP167:HE1 - B:VAL275:O | 2.10242 | Hydrogen Bond | Conventional H-Bond |
| B:THR276:HG1 - A:GLN70:OE1 | 2.49007 | Hydrogen Bond | Conventional H-Bond |
| B:GLY278:CA - A:ASN66:OD1 | 3.62 | Hydrogen Bond | Carbon H-Bond |
| B:VAL275:N - A:TYR7 | 4.18541 | Electrostatic | Pi-Cation |
| B:VAL275:N - A:TRP167 | 4.12957 | Electrostatic | Pi-Cation |
| B:VAL275:HT3 - A:TRP167 | 3.11323 | Hydrogen Bond;Electrostatic | Pi-Cation;Pi-Donor H-Bond |
| A:ASP77:OD1 - B:TYR283 | 3.26765 | Electrostatic | Pi-Anion |
| A:TRP147 - B:TYR283 | 5.19777 | Hydrophobic | Pi-Pi T-shaped |
| A:TRP147 - B:TYR283 | 4.86133 | Hydrophobic | Pi-Pi T-shaped |
| B:THR276:C,O;TRP277:N - A:TYR159 | 4.43766 | Hydrophobic | Amide-Pi Stacked |
| A:VAL67 - B:VAL275 | 4.9401 | Hydrophobic | Alkyl |
| A:ALA150 - B:PRO282 | 4.38864 | Hydrophobic | Alkyl |
| A:ALA152 - B:PRO282 | 4.51101 | Hydrophobic | Alkyl |
| A:TYR7 - B:VAL275 | 4.84813 | Hydrophobic | Pi-Alkyl |
| B:TRP277 - A:ARG163 | 4.97154 | Hydrophobic | Pi-Alkyl |
| B:TYR283 - A:LYS146 | 4.866 | Hydrophobic | Pi-Alkyl |

**Supplementary Table 15:** Interaction of predicted CD4+ T-cell epitope “DNFNVYKATRPYLAH” in CHIKV E2 with MHC class-II alleles having affinity IC50 < 100.

| **Name** | **Distance** | **Category** | **Types** |
| --- | --- | --- | --- |
| A:ARG74:HH12 - B:HIS194:OXT | 2.03445 | Hydrogen Bond; Electrostatic | Salt Bridge;  Attractive Charge |
| B:ARG189:HH11 - A:GLU9:OE1 | 2.60754 | Hydrogen Bond; Electrostatic | Salt Bridge;  Attractive Charge |
| B:ARG189:HH21 - A:GLU139:OE2 | 2.15812 | Hydrogen Bond; Electrostatic | Salt Bridge;  Attractive Charge |
| B:ARG189:HH22 - A:GLU139:OE1 | 3.19275 | Hydrogen Bond; Electrostatic | Salt Bridge;  Attractive Charge |
| A:ARG74:NH2 - B:HIS194:O | 5.40345 | Electrostatic | Attractive Charge |
| B:ARG189:NH1 - A:GLU139:OE1 | 5.40197 | Electrostatic | Attractive Charge |
| A:GLN7:HE21 - B:THR188:O | 2.10325 | Hydrogen Bond | Conventional H-Bond |
| A:GLN7:HE22 - B:ALA187:O | 2.50333 | Hydrogen Bond | Conventional H-Bond |
| B:ASN181:HN - B:ASP180:OD2 | 3.02643 | Hydrogen Bond | Conventional H-Bond |
| B:ASN181:HD22 - B:ASN183:O | 2.41857 | Hydrogen Bond | Conventional H-Bond |
| B:ASN183:HD22 - B:ASN181:O | 2.56684 | Hydrogen Bond | Conventional H-Bond |
| B:ARG189:HE - A:GLN7:O | 2.01015 | Hydrogen Bond | Conventional H-Bond |
| B:ARG189:HH11 - B:PRO190:O | 2.81933 | Hydrogen Bond | Conventional H-Bond |
| B:TYR191:HH - A:ASN60:OD1 | 2.62608 | Hydrogen Bond | Conventional H-Bond |
| B:TYR191:HH - A:ASN60:O | 2.76261 | Hydrogen Bond | Conventional H-Bond |
| B:LEU192:HN - A:ASP64:OD1 | 2.6713 | Hydrogen Bond | Conventional H-Bond |
| A:PHE46:CA - B:ASP180:OD2 | 3.75997 | Hydrogen Bond | Carbon H-Bond |
| B:THR188:CA - A:ASN60:OD1 | 3.76904 | Hydrogen Bond | Carbon H-Bond |
| A:GLU9:OE1 - B:TYR191 | 4.06679 | Electrostatic | Pi-Anion |
| B:ASP180:OD1 - A:PHE46 | 3.83355 | Electrostatic | Pi-Anion |
| B:ASP180:OD2 - B:PHE182 | 4.92297 | Electrostatic | Pi-Anion |
| A:VAL63:CG1 - B:TYR191 | 3.91953 | Hydrophobic | Pi-Sigma |
| B:ASP180:CB - B:PHE182 | 3.89511 | Hydrophobic | Pi-Sigma |
| A:PHE49 - B:PHE182 | 4.90337 | Hydrophobic | Pi-Pi T-shaped |
| B:ALA193 - A:MET71 | 4.45324 | Hydrophobic | Alkyl |
| A:PHE52 - B:ALA187 | 4.90028 | Hydrophobic | Pi-Alkyl |
| A:HIS141 - B:LEU192 | 5.09699 | Hydrophobic | Pi-Alkyl |
| B:TYR185 - A:ILE29 | 5.20455 | Hydrophobic | Pi-Alkyl |
| B:TYR185 - A:ALA50 | 5.19016 | Hydrophobic | Pi-Alkyl |

**Supplementary Table 16:** Interaction of predicted CD4+ T-cell epitope “RAGLFVRTSAPCTIT” in CHIKV E2 with MHC class-II alleles having affinity IC50 < 100.

| **Name** | **Distance** | **Category** | **Types** |
| --- | --- | --- | --- |
| A:LYS73:HZ1 - B:THR194:OXT | 1.88765 | Hydrogen Bond;  Electrostatic | Salt Bridge;  Attractive Charge |
| A:GLN7:HE21 - B:ARG186:O | 2.79548 | Hydrogen Bond | Conventional H-Bond |
| A:ASN60:HD21 - B:ARG186:O | 2.31115 | Hydrogen Bond | Conventional H-Bond |
| B:ARG180:HH12 - A:SER51:O | 2.13953 | Hydrogen Bond | Conventional H-Bond |
| B:ARG180:HH22 - A:SER51:O | 2.18784 | Hydrogen Bond | Conventional H-Bond |
| B:GLY182:HN - A:GLU1:OE2 | 2.07076 | Hydrogen Bond | Conventional H-Bond |
| B:LEU183:HN - B:ALA181:O | 1.97795 | Hydrogen Bond | Conventional H-Bond |
| B:ARG186:HE - A:GLN7:OE1 | 1.91948 | Hydrogen Bond | Conventional H-Bond |
| B:ARG186:HH21 - A:GLN7:OE1 | 2.76271 | Hydrogen Bond | Conventional H-Bond |
| B:SER188:HN - A:ASN60:OD1 | 1.99184 | Hydrogen Bond | Conventional H-Bond |
| B:SER188:HG - A:GLU9:OE2 | 2.15927 | Hydrogen Bond | Conventional H-Bond |
| B:THR187:CA - A:ASN60:OD1 | 3.17128 | Hydrogen Bond | Carbon H-Bond |
| B:ARG186:NH2 - B:PHE184 | 4.34845 | Electrostatic | Pi-Cation |
| A:ILE5 - B:LEU183 | 5.47613 | Hydrophobic | Alkyl |
| A:ALA50 - B:ARG180 | 4.0315 | Hydrophobic | Alkyl |
| A:ALA50 - B:LEU183 | 4.55289 | Hydrophobic | Alkyl |
| A:VAL63 - B:PRO190 | 4.52815 | Hydrophobic | Alkyl |
| A:ALA66 - B:ILE193 | 4.68615 | Hydrophobic | Alkyl |
| B:ALA181 - B:LEU183 | 5.08719 | Hydrophobic | Alkyl |
| A:PHE22 - B:LEU183 | 5.19664 | Hydrophobic | Pi-Alkyl |
| A:PHE46 - B:ALA181 | 5.09856 | Hydrophobic | Pi-Alkyl |

**Supplementary Table 17:** Interaction of predicted CD4+ T-cell epitope “YYYELYPTMTVVVVS” with CHIKV E2 MHC class-II alleles having affinity IC50 < 100.

| **Name** | **Distance** | **Category** | **Types** |
| --- | --- | --- | --- |
| A:GLU1:HT1 - B:TYR180:OH | 2.00921 | Hydrogen Bond | Conventional H-Bond |
| A:GLN7:HE21 - B:MET188:O | 2.08716 | Hydrogen Bond | Conventional H-Bond |
| A:ASN60:HD21 - B:MET188:O | 2.52794 | Hydrogen Bond | Conventional H-Bond |
| B:TYR180:HH - A:HIS3:O | 2.85725 | Hydrogen Bond | Conventional H-Bond |
| B:GLU183:HN - B:TYR180:O | 2.22548 | Hydrogen Bond | Conventional H-Bond |
| B:LEU184:HN - B:TYR180:O | 2.09954 | Hydrogen Bond | Conventional H-Bond |
| B:THR187:HG1 - A:GLN7:OE1 | 2.13758 | Hydrogen Bond | Conventional H-Bond |
| B:THR187:HG1 - B:THR187:O | 2.30144 | Hydrogen Bond | Conventional H-Bond |
| B:VAL190:HN - A:GLU9:OE2 | 2.25832 | Hydrogen Bond | Conventional H-Bond |
| B:VAL193:HN - A:GLU9:O | 1.95007 | Hydrogen Bond | Conventional H-Bond |
| B:SER194:HG - A:TYR11:O | 2.03741 | Hydrogen Bond | Conventional H-Bond |
| B:THR189:CA - A:GLU9:OE2 | 3.37801 | Hydrogen Bond | Carbon H-Bond |
| A:MET71:SD - B:SER194:OXT | 3.23192 | Other | Sulfur-X |
| B:MET188:SD - A:PHE52 | 5.21741 | Other | Pi-Sulfur |
| A:HIS3 - B:TYR180 | 4.78875 | Hydrophobic | Pi-Pi Stacked |
| A:PHE22 - B:TYR185 | 4.70906 | Hydrophobic | Pi-Pi Stacked |
| A:PHE46 - B:TYR181 | 5.41283 | Hydrophobic | Pi-Pi Stacked |
| A:PHE24 - B:TYR181 | 5.04488 | Hydrophobic | Pi-Pi T-shaped |
| A:HIS141 - B:VAL193 | 5.01245 | Hydrophobic | Pi-Alkyl |
| A:PHE143 - B:VAL193 | 5.08865 | Hydrophobic | Pi-Alkyl |
| B:TYR180 - B:PRO186 | 5.26196 | Hydrophobic | Pi-Alkyl |
| B:TYR185 - A:ILE29 | 5.37059 | Hydrophobic | Pi-Alkyl |
| B:TYR185 - A:ALA50 | 5.49387 | Hydrophobic | Pi-Alkyl |

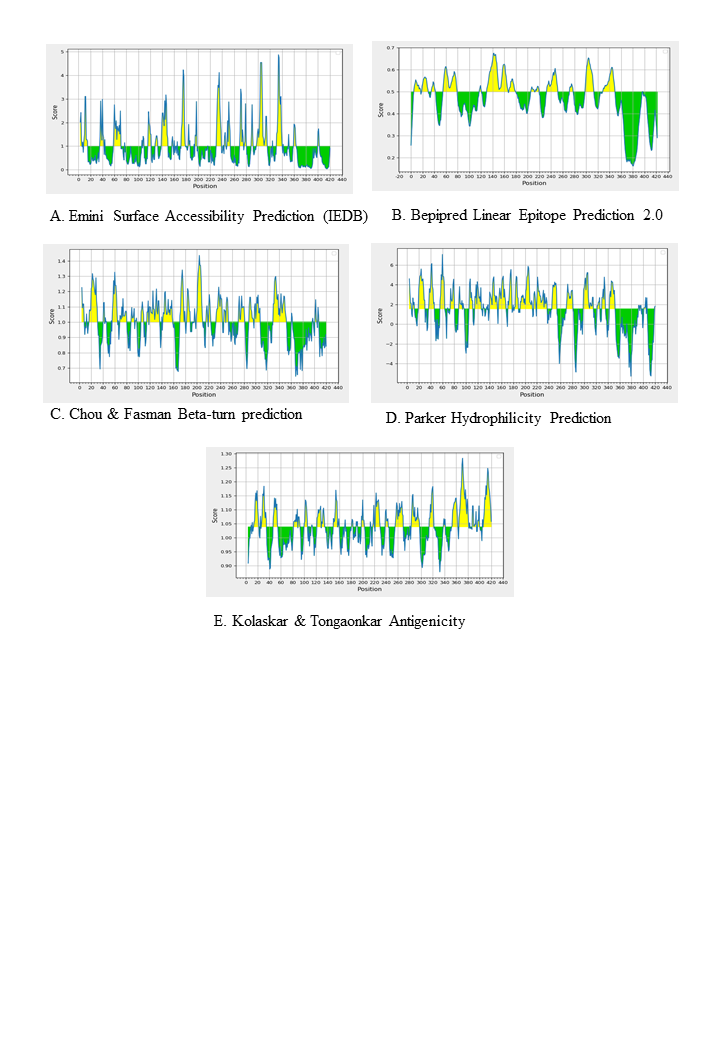

**Supplementary Figure 5 (A-E):** Many epitope peptides from CHIKV E2 were predicted by various tools: **A.** Emini Surface Accessibility Prediction (IEDB). **B.** Bepipred Linear Epitope Prediction 2.0. **C.** Chou & Fasman Beta-turn prediction. **D.** Kolaskar & Tongaonkar Antigenicity prediction. **E.** Parker Hydrophilicity Prediction.

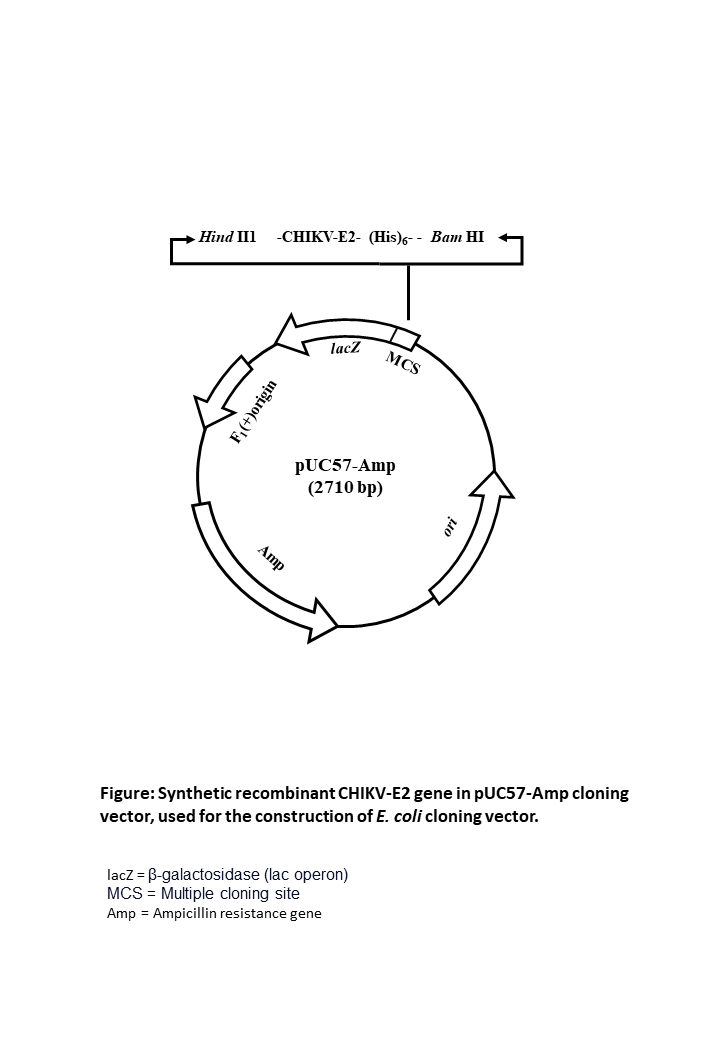

**Supplementary Figure 6:** Synthetic recombinant CHIKV-E2 gene in pUC57-Amp cloning vector, used for the construction of E. coli cloning vector

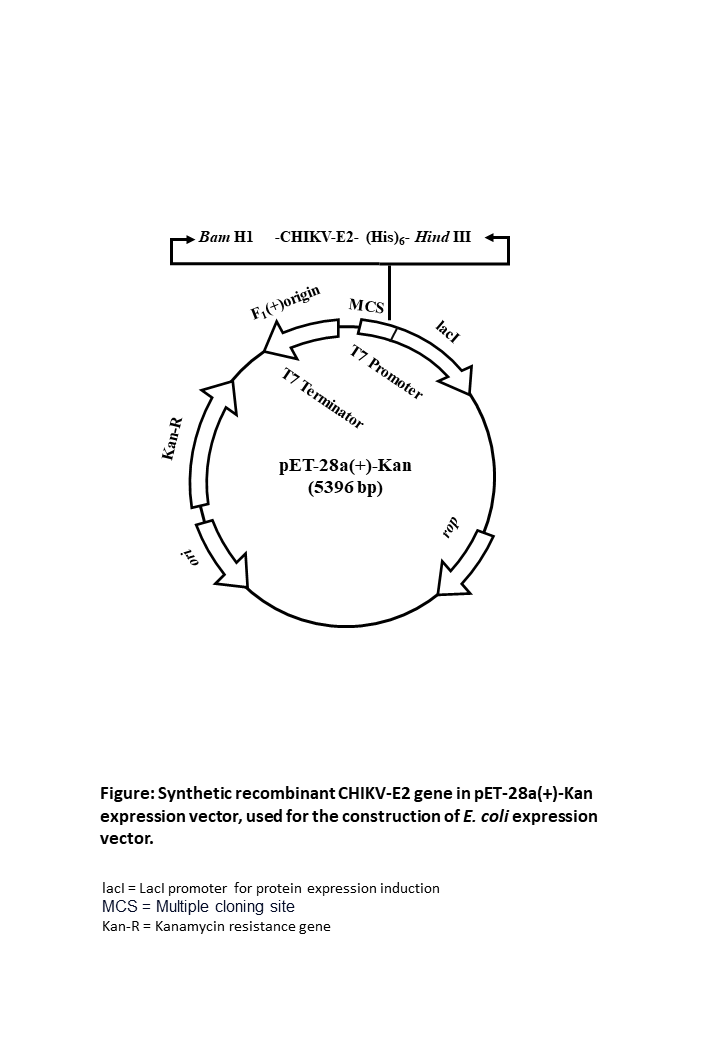

**Supplementary Figure 7:** Synthetic recombinant CHIKV-E2 gene in pET-28a(+)-Kan expression vector, used for the construction of E. coli expression vector.

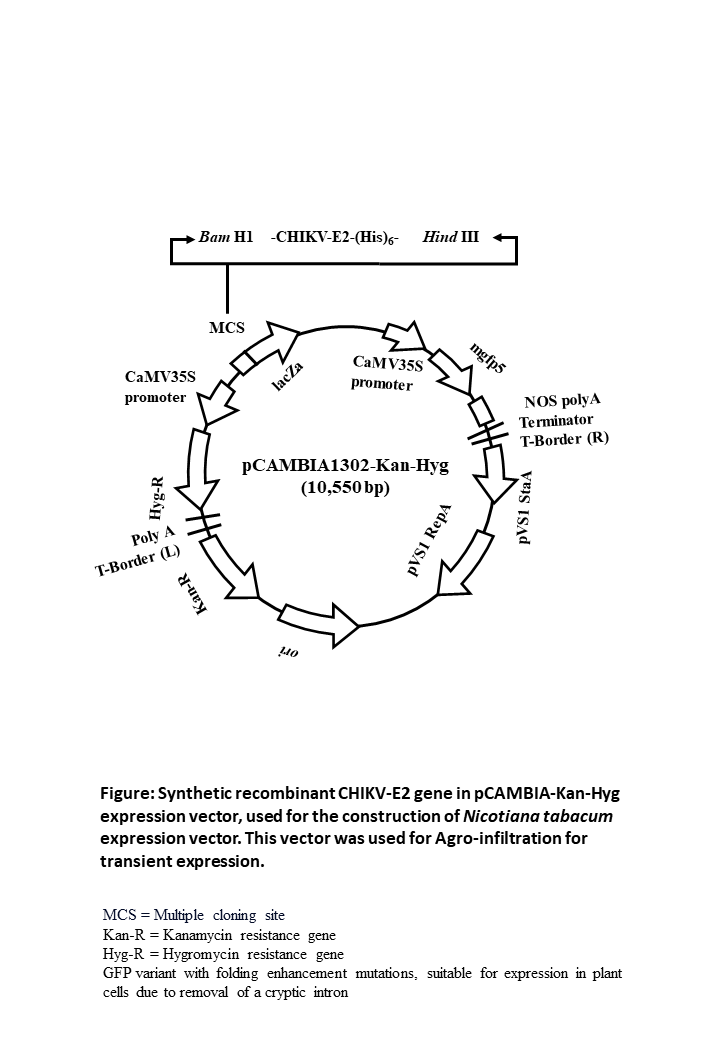

**Supplementary Figure 8:** Synthetic recombinant CHIKV-E2 gene in pCAMBIA-Kan-Hyg expression vector, used for the construction of *Nicotiana tabacum* expression vector. This vector was used for Agro-infiltration for transient expression.

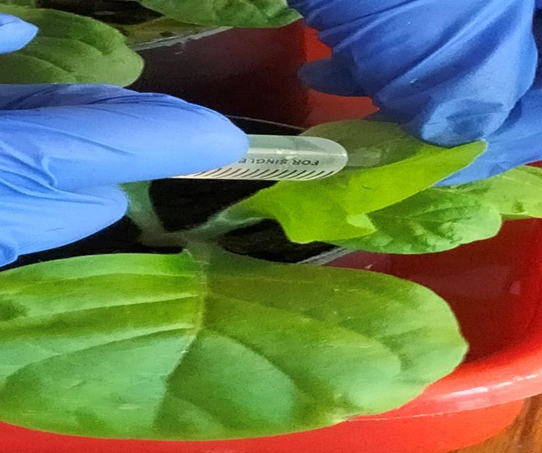

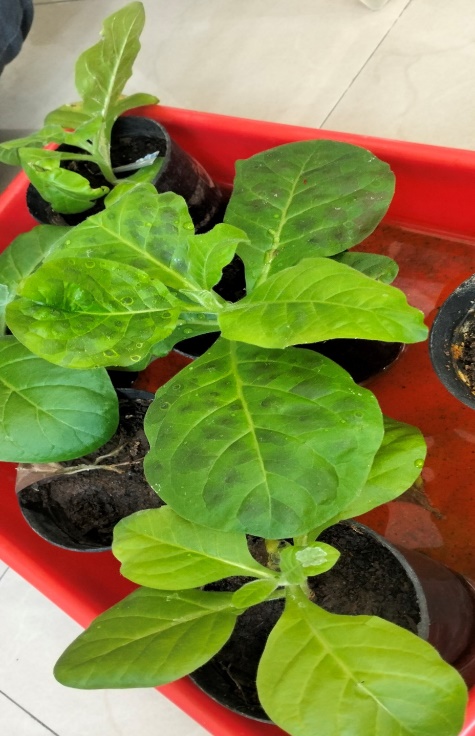

**Supplementary Figure 9:** Transient transformation of tobacco leaf through the abaxial surface using transformed *A. tumefaciens.*
